# Elucidation of global and local genomic epidemiology of *Salmonella enterica* serovar Enteritidis through multilevel genome typing

**DOI:** 10.1101/2020.06.30.169953

**Authors:** Lijuan Luo, Michael Payne, Sandeep Kaur, Dalong Hu, Liam Cheney, Sophie Octavia, Qinning Wang, Mark M. Tanaka, Vitali Sintchenko, Ruiting Lan

## Abstract

*Salmonella enterica* serovar Enteritidis is a major cause of foodborne *Salmonella* infections and outbreaks in humans. Effective surveillance and timely outbreak detection are essential for public health control. Multilevel genome typing (MGT) with multiple levels of resolution has been previously demonstrated as a promising tool for this purpose. In this study, we developed MGT with nine levels for *S.* Enteritidis and characterised the genomic epidemiology of *S.* Enteritidis in detail. We examined 26,670 publicly available *S*. Enteritidis genome sequences from isolates spanning 101 years from 86 countries to reveal their spatial and temporal distributions. Using the lower resolution MGT levels, globally prevalent and regionally restricted sequence types (STs) were identified; avian associated MGT4-STs were found that were common in human cases in the USA were identified; temporal trends were observed in the UK with MGT5-STs from 2014 to 2018, revealing both long lived endemic STs and the rapid expansion of new STs. Using MGT3 to MGT6, we identified MDR associated STs at various MGT levels, which improves precision of detection and global tracking of MDR clones. We also found that the majority of the global *S*. Enteritidis population fell within two predominant lineages, which had significantly different propensity of causing large scale outbreaks. An online open MGT database has been established for unified international surveillance of *S*. Enteritidis. We demonstrated that MGT provides a flexible and high-resolution genome typing tool for *S*. Enteritidis surveillance and outbreak detection.

**Impact statement:** *Salmonella enterica* serovar Enteritidis is a common foodborne pathogen that can cause large outbreaks. Surveillance and high-resolution typing are essential for outbreak prevention and control. Genome sequencing offers unprecedented power for these purposes and a standardised method or platform for the interpretation, comparison and communication of genomic typing data is highly desirable. In this work, we developed a genomic typing scheme called Multilevel Genome Typing (MGT) for *S*. Enteritidis. We analysed 26,670 publicly available genomes of *S.* Enteritidis using MGT. We characterised the geographic and temporal distribution of S. Enteritidis MGT types as well as their association with multidrug resistance (MDR) and virulence genes. A publicly available MGT database for *S*. Enteritidis was established, which has the potential facilitate the unified global public health surveillance for this pathogen.

**Data Summary:**

1. The MGT database for *S.* Enteritidis is available at https://mgtdb.unsw.edu.au/enteritidis/.
2. All accession numbers of the public available genomes were available in the MGT database and Data Set S1, Tab 1. And there were no newly sequenced data in this study.
3. Supplementary material: Supplementary Fig. S1 to S7, supplementary methods and supporting results about the evaluation of potential repeat sequencing bias.
4. Data Set S1: Supporting tables of the main results.
5. Data Set S2. Supporting tables of the repeat sequencing bias evaluation by removing the potential repeat sequencing isolates. Note outbreak isolates may also be removed.

## Introduction

Nontyphoidal *Salmonella* is ranked second in foodborne disease in Europe and North America [1, 2], with *Salmonella enterica* serovar Enteritidis a dominant serovar in many countries [3, 4]. *S.* Enteritidis mainly causes human gastrointestinal infections leading to diarrhoea. However, invasive infections manifesting as sepsis, meningitis and pneumonia, have recently been reported [5, 6]. As many *S.* Enteritidis strains causing invasive infections carry plasmids which confer multidrug resistance (MDR), the spread of these strains internationally is a major threat to global health [7, 8]. Additionally, *S.* Enteritidis has caused numerous large-scale national and international outbreaks with complex transmission pathways [3, 9, 10]. While there have been limited studies of the total global epidemiology of *S.* Enteritidis genomic subtypes [5, 11], the geographic distribution, outbreak propensity, MDR and evolutionary characteristics of different lineages of *S.* Enteritidis have not been systematically evaluated using the large numbers of newly available public genome sequences. A rapid, stable and standardized global genomic typing strategy for *S.* Enteritidis is required for the high-resolution and scalable surveillance of outbreaks, tracking of international spread and MDR profiles of *S.* Enteritidis.

Sequence types (STs) are based on exact matching of genes between isolates. The well-established seven gene multi-locus sequence typing (MLST) of *Salmonella* is widely used [12], while the vast majority of *S.* Enteritidis are assigned to ST11, as only seven housekeeping genes are compared. STs of higher resolution are required. With this in mind core genome MLST (cgMLST) and whole genome MLST (wgMLST) schemes of *Salmonella* were developed including 3002 and 21,065 loci, respectively [13]. However, STs based on cgMLST and wgMLST are too diverse to offer useful relatedness information at anything but the finest scales. This issue was addressed by single linkage hierarchic typing systems, i.e., Single Nucleotide Polymorphism (SNP) address and Hierarchical Clustering of cgMLST (HierCC) [14–16]. SNP address is based on the single linkage hierarchic clustering method, which allows for different SNP differences 250, 100, 50, 25, 10, 5 and 0 SNPs [15]. These methods had been successfully used in outbreak tracing and epidemiology [14, 17–19]. However, one major disadvantage of the pairwise comparison based single linkage clustering method is that the cluster types called may merge when additional data is added and differences in order of addition can result in different clusters [20]. For example, when an isolate meets the SNP/allele difference threshold to two clusters, this isolate would act as a bridge connecting the two clusters and merging them into a single cluster. This scenario will almost certainly occur in a large database with ever expanding numbers of sequences, especially at finer resolutions, and would be an obstacle for long term epidemiological investigation and comparison [20]. We have recently developed a novel core genome based method called multilevel genome typing (MGT), an exact matching method to assign multiple resolutions of relatedness without the need for clustering [20]. MGT is a genome sequence based typing system, including multiple MLST schemes with increasing resolution from the classic seven gene MLST to cgMLST where each scheme is used to independently assign an ST. An MGT scheme and a public database were established for *Salmonella enterica* serovar Typhimurium where seven to 5268 loci were included in MGT1 to 9 [20]. MGT1 corresponds to the classic seven gene MLST scheme of *S. enterica* [12]. For *S.* Typhimurium, the stable STs from MGT levels of appropriate resolution were found to be useful in identifying DT104, the MDR lineage, as well as African invasive lineages [20]. MGT of *S*. Typhimurium was also shown to perform well in outbreak identification and source tracing [20].

Here, we describe an MGT scheme and public database for *S.* Enteritidis. By using MGT, we systematically examine the global and local epidemiology of *S.* Enteritidis over time, and evaluate the application of this scheme to the outbreak evaluation and the monitoring of multidrug resistant STs.

## Methods

### S. Enteritidis sequences used in this study

Raw reads of 26,670 *S.* Enteritidis whole genome sequences (WGS) downloaded from the European Nucleotide Archive (ENA) passed the quality filtering criteria by Quast [21, 22] (**Data Set S1, Tab 1)**. The epidemiological metadata of those isolates were collated from NCBI and EnteroBase [13]. Trimmomatic v0.39 was used with default settings to trim raw reads [23]. The trimmed raw reads were then assembled using either SPAdes v3.13.0 for core genome definition or SKESA v2.3 for MGT typing [24, 25]. Quality assessment for the assemblies were performed using Quast v5.0.2 [21]. Thresholds of the assembly quality were in accordance with Robertson *et al*’s criteria [22]. Kraken v0.10.5 was used to identify contamination in the assembled genomic sequences or problematic taxonomic identification [26]. Seqsero v1.0 and SISTR v1.0.2 were used to verify serotype assignments [27, 28].

### Core gene definition of *S.* Enteritidis

To define the core genome of *S.* Enteritidis, a total of 1801 genomes were selected based on the ribosomal sequence type [13] and strain metadata. The trimmed raw reads were then assembled using SPAdes v3.13.0 [24]. The assemblies were annotated with Prokka v1.12[29]. The core genes and core intergenic regions were defined with Roary v3.11.2 and Piggy v3.11.2, respectively (Fig. S1). A total of 113 sRNA regions of SL1344 were searched in the 1054 intergenic regions using BLAST, with identity and coverage threshold of 70% and 60%, respectively [30]. Both SeqSero v1.0 and SISTR v1.0.2 were used to verify whether the isolates were S. Enteritidis [27, 28]. The detailed methods for isolates selection and core genome identification were described in Supplementary material.

### Establishment of MGT scheme for *S.* Enteritidis and MGT allele calling

The initial allele sequence of each locus used for MGT were extracted from the complete genome sequence of *S.* Enteritidis strain P125109 (Genbank accession NC_011294.1). The first eight levels of MGT were identical to the MGT scheme for *S.* Typhimurium [20], in which MGT1 refers to the classical seven gene MLST for the *Salmonella* genus [12]. Loci used for levels MGT2 to MGT8 were orthologous to those in the *S.* Typhimurium MGT with the exception of one gene which was removed at MGT8 because of a paralogue in *S.* Enteritidis. MGT9 contains all loci in MGT8 as well as the core genes and core intergenic regions of *S.* Enteritidis.

The raw read processing and genome assembly procedures were the same as above, except that SKESA v2.3 was used to assemble the raw reads and only SISTR v1.0.2 was used for the molecular serotype identification [25, 28]. SKESA v2.3 offers higher per base accuracy and speed than SPADES and was therefore used in the allele calling pipeline [25]. For the assemblies that passed quality control by the criteria of Robertson *et al.* [22], a custom script was used to assign alleles, STs and clonal complexes (CCs) to those isolates at levels from MGT2 to MGT9 [20]. Allele, ST, CC and outbreak detection cluster (ODC) assignments were performed as previously described [20]. Briefly, if one or more allele difference was observed, a new ST was assigned to the isolate. The same strategy was also used for assigning a new CC, if at least two allele differences were observed comparing to any of STs in existing CCs in the database [20]. ODC is an extension of the CC clustering method, but it was performed only at MGT9 level [20]. For example, any two MGT9 STs with no more than 2, 5, or 10 allele differences were assigned the same ODC2, ODC5 or ODC10 type [20].

### Geographic distribution and temporal variation analysis

We first determined which MGT level would be the most useful in describing the global epidemiology of *S.* Enteritidis by summarizing STs that contained at least 10 isolates for their distribution across continents. STs where more than 70% of isolates were from a single continent were identified at each level of MGT (Fig. 2a). MGT4 STs matching this criterion had the highest percentage of all the levels at 76.3% of the 26,670 isolates, thus MGT4 was chosen to describe the continental distribution of *S*. Enteritidis. The USA states variation map and the UK monthly variation column chart were produced using Tableau v2019.2 [31].

### Antibiotic resistance gene and plasmid specific gene identification

Abricate v1.0.1 (https://github.com/tseemann/abricate) was used for identification of antibiotic resistance genes with the ResFinder v3.0 database [32], and plasmid specific genes with the PlasmidFinder v2.1 database [33]. The cut-off of the presence of a gene was set as both identity and coverage >= 90% [32]. At each MGT level, STs (>=10 isolates each) with more than 80% of isolates harbouring antibiotic resistance genes, were collected from MGT2 to MGT9 and were defined as high-antibiotic-resistant STs. The STs at each level of MDR were non-redundant meaning that no isolate was counted more than once.

### Phylogenetic analysis

A phylogenetic tree was constructed using Parsnp v1.2, which called core-genome SNPs [34, 35]. Potential recombinant SNPs were removed using both Gubbins v2.0.0 and Recdetect v6.0 [36, 37]. Beast v1.10.4 was then used to estimate the mutation rate based on mutational SNPs [38]. A total of 24 combinations of clock and population size models were evaluated with the MCMC chain of 100 million states. Tracer v1.7.1 was used to identify the optimal model and to estimate population expansion over time [39]. The detailed methods for phylogenetic analysis were described in **Supplementary material.**

### Virulence genes and *Salmonella* pathogenicity islands (SPIs) distribution in the *S.* Enteritidis population

We compared the presence of virulence determinants in the main lineages of *S*. Enteritidis. Virulence genes from the Virulence Factor Database (VFDB) were identified in all the *S.* Enteritidis genomes using Abricate v1.0.1 (https://github.com/tseemann/abricate), with identity threshold of 70% and coverage of 50% [40]. Using the same blast threshold, a total of 23 reported SPIs from SPI-1 to SPI-23 were also identified in all the *S.* Enteritidis genomes [41].

### Evaluation of the effect of repeat sequencing on the dataset for epidemiological analysis

To evaluate any bias that may be caused by resequencing of the same isolate, we identified all isolates of the same ST based on MGT9 and same metadata based on collection country, collection year and month, and source type. Such isolates were conservatively treated as repeat sequencing of the same isolate and such “duplicates” were removed from the dataset. We re-analysed the reduced dataset and compared against the original dataset by calculating Kendall’s tau [42]. The detailed results are shown in Supplementary material and Data Set S2, Tab 1 to Tab 6.

## Results

### Establishing the MGT system for *S.* Enteritidis

The core genome of *S.* Enteritidis was defined at >=96% identity and presence rate of >=99% of the 1801 sampled isolates (Fig. S1). The core genome of *S.* Enteritidis included 3932 genes, with 977 not found in the *Salmonella enterica* core as well as 1054 *S.* Enteritidis core intergenic regions (Fig. 1a) (Data Set S1, Tab 2). We also searched for the presence of small RNA (sRNA) in the intergenic regions with 37 (32.7%, 37/113) sRNAs observed in 36 (3.4%, 36/1054) intergenic regions.

**Fig. 1.**
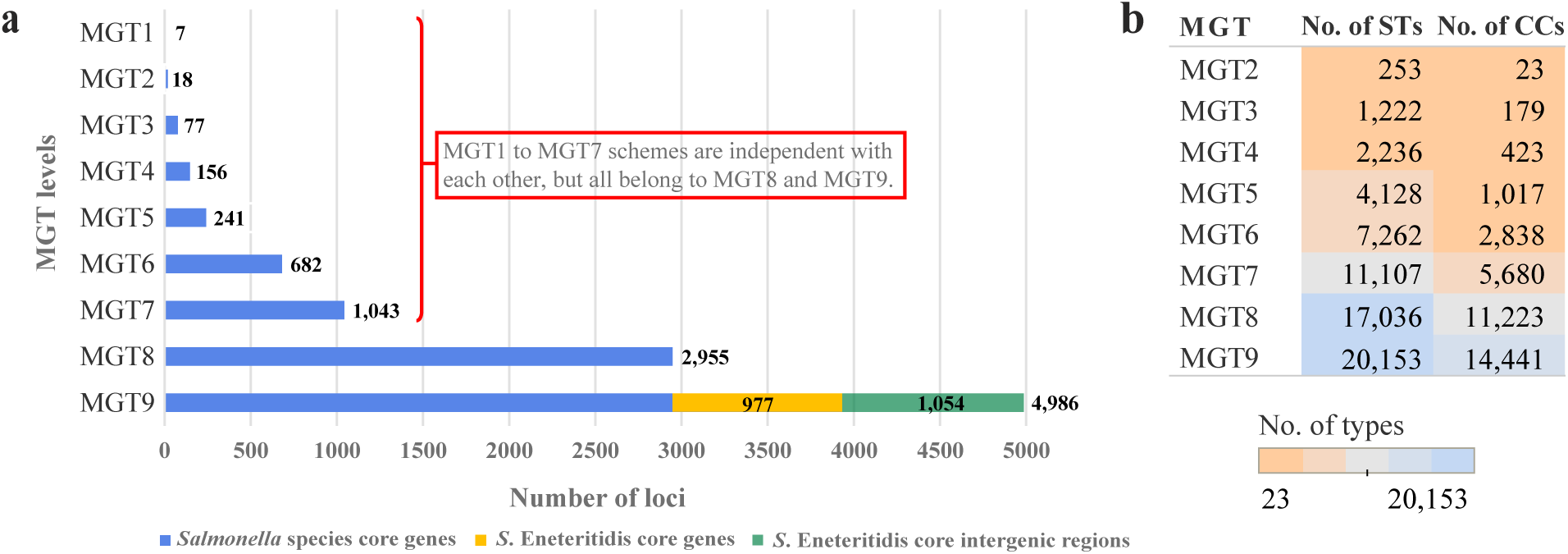
Makeup and assignments of each *S.* Enteritidis MGT level. **a.** Number of loci included in each MGT level. The first eight levels are composed of *Salmonella enterica* core genes, which are orthologous to those of the *S.* Typhimurium MGT1 to MGT8, except for one gene at MGT8 which was excluded due to duplication in *S.* Enteritidis. The MGT9 scheme includes core genes of S. Enteritidis (those not belonging to the *Salmonella* core coloured in yellow) and core intergenic regions (coloured in green). The number behind or within each bar refers to the number of loci included. **b.** The number of sequence types (ST) and clonal complexes (CC) assigned at each MGT level among the 26,670 genomes analysed. CC includes STs of no more than one allele difference. The numbers refer to the number of STs or CCs types assigned at each level, which were also indicated by colour. As MGT9 included 4986 loci with highest subtyping resolution, the 26,670 isolates were subtyped into 20,153 different ST types and 14,441 CC clustering types at MGT9.

The first eight schemes of *S.* Enteritidis MGT used the same 2955 core genes of *Salmonella enterica* as the MGT of *S.* Typhimurium, and the rationale of locus selection has been fully described by Payne *et al* [20]. MGT1 corresponded to the classic seven gene MLST of *Salmonella* [12]. A total of 4986 loci were incorporated in the MGT9 scheme, including the 3932 *S.* Enteritidis core genes and 1054 core intergenic regions defined (Fig. 1a).

A total of 26,670 *S.* Enteritidis genomes with publicly available raw reads were analysed using the MGT scheme. These publicly available genomes were collected from 26 source types with 49.9% of isolates collected from humans and 8.9% from avian sources; they were collected from 86 countries with the majority of the isolates from the United States (47.3%) and United Kingdom (35.9%); and they were collected between the years 1917 and 2018 with 2014 (11.5%), 2015 (14.6%), 2016 (14.0%), 2017 (12.8%) and 2018 (3.7%) having more than 500 isolates each (Fig. S2a). At each MGT level, each isolate was assigned an ST and a clonal complex (CC). A CC in this study was defined as a group of STs with one allele difference [12, 43]. The number of STs and CCs at each MGT level is shown in Fig. 1b. As the resolution of typing increased from MGT1 to MGT9, an increasing number of STs and CCs were assigned. At MGT2, the 26,670 isolates were subtyped into 252 STs and 23 CCs. By contrast, MGT9 divided the isolates into 20,153 different STs and 14,441 CCs. The MGT scheme of *S.* Enteritidis is available through a public online database (https://mgtdb.unsw.edu.au/enteritidis/).

### The international or global epidemiology of *S.* Enteritidis by MGT

Of the 26,670 isolates typed by MGT, 25,207 have country metadata. By the 7-gene MLST (or MGT1) scheme, ST11 was the dominant type representing 94.8% of the isolates, followed by ST183 representing 1.6%. ST183 is an endemic ST prevalent in Europe and was divided into two main phage types, PT11 and PT66 [44]. Using MGT, ST183 (or MGT1-ST183) can be divided into seven STs at MGT2 level. Interestingly, 97% (137/141) of the PT11 isolates belonged to MGT2-ST3, and 100% (24/24) of the PT66 belonged to MGT2-ST82, highlighting the potential for backward compatibility of MGT with traditional typing data.

For the predominant ST11 isolates (or MGT1-ST11), we found that the optimal MGT level to describe their global epidemiology was MGT4, based on the distribution of each ST in different continents (Fig. 2a). At the MGT4 level, the 26,670 isolates were subtyped into 2,236 STs and 423 CCs. Among the 2,236 MGT4-STs, 163 STs were predominantly found in only one continent (defined as ≥70% from one continent). These 163 STs contained 20,341 of the 26,670 genomes (76.3%).

**Fig. 2.**
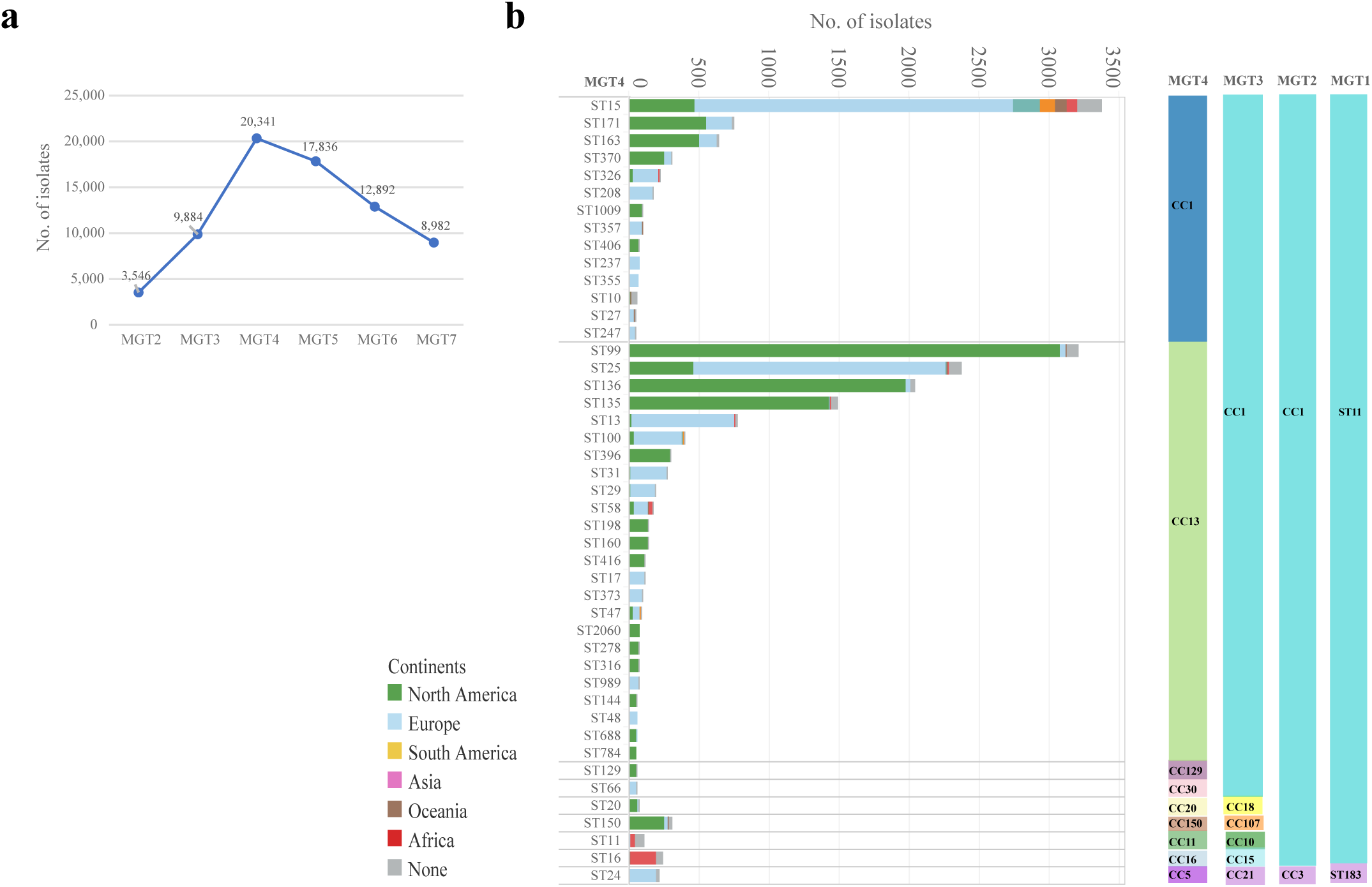
MGT4 STs offer a useful description of continent specific and global clades. **a.** At each MGT level from MGT2 to MGT8, we identified continent restricted STs where more than 70% of the isolates belonged to the same continent. MGT4 was found to have the highest number of isolates belonged to these continental limited STs. **b.** There were 45 MGT4-STs of more than 50 isolates representing 73.5% of the total 26670 isolates, and the majority of these STs belonged to MGT4-CC1 and CC13. The number, size and continental makeup of these STs is shown with the left Y axis showing MGT4 STs, the right Y axis showing MGT4 CCs and the X axis showing number of isolates, continental distribution of each ST is shown by different colours in each row.

The distribution of STs varied between continents. The following MGT4-CC1 STs, ST171, ST163, ST370, ST1009, and MGT4-CC13 STs, ST99, ST136, ST135, ST396, ST198, ST160, ST416 were the most prevalent STs in North America (Fig. 2b). While in Europe, the following MGT4-CC1 STs, ST15, ST208, ST357, ST237 and MGT4-CC13 STs, ST25, ST13, ST100, ST31, ST29 were common. However, the North America and Europe *S.* Enteritidis sequences analysed were mostly obtained from the USA and UK, which represented 94.6% and 92.5% of isolates from these two continents (Fig. S2a). In Africa, MGT4-ST11 (also described by MGT3-ST10) was more prevalent in West Africa and MGT4-ST16 (also described by MGT3-ST15) in Central/Eastern Africa (Fig. S3). Finally, MGT4-ST15 was a global ST which was observed in all continents (Fig. 2b).

These dominant STs can be further grouped into CCs which offered a more inclusive picture. Among the 423 MGT4-CCs at MGT4, 10 represented 94.1% of all the isolates. The top two CCs, MGT4-CC1 and MGT4-CC13 accounted for 88.0% of the isolates (Fig. 2b). MGT4-CC1 was prevalent in all six continents, while MGT4-CC13 was more common in North America and Europe (Fig. 2b, Fig. S4).

### The national or local epidemiology of *S.* Enteritidis by MGT

As the majority of the *S.* Enteritidis sequences analysed in this study were from the USA and UK, we compared the distribution of the STs between these two countries. We used MGT4 and MGT5 levels to describe the data. A total of 39 MGT4 STs with more than 50 isolates represented 75.1% of the USA and UK isolates and each country had its own specific types (Fig. 3a). MGT4-ST99, ST136, ST135, ST171, ST163, ST370, ST396 and ST150 were the main STs in the USA, whereas MGT4-ST15, ST25, ST13, ST100, ST31, ST326, ST24, ST208, ST29 were the main STs in the UK. MGT4-ST15 and ST25 were further subtyped into 34 MGT5-STs (each with more than 20 isolates), of which the majority were mainly observed in the UK, except for MGT5-ST412 and ST387 in the USA (Fig. 3b).

**Fig. 3.**
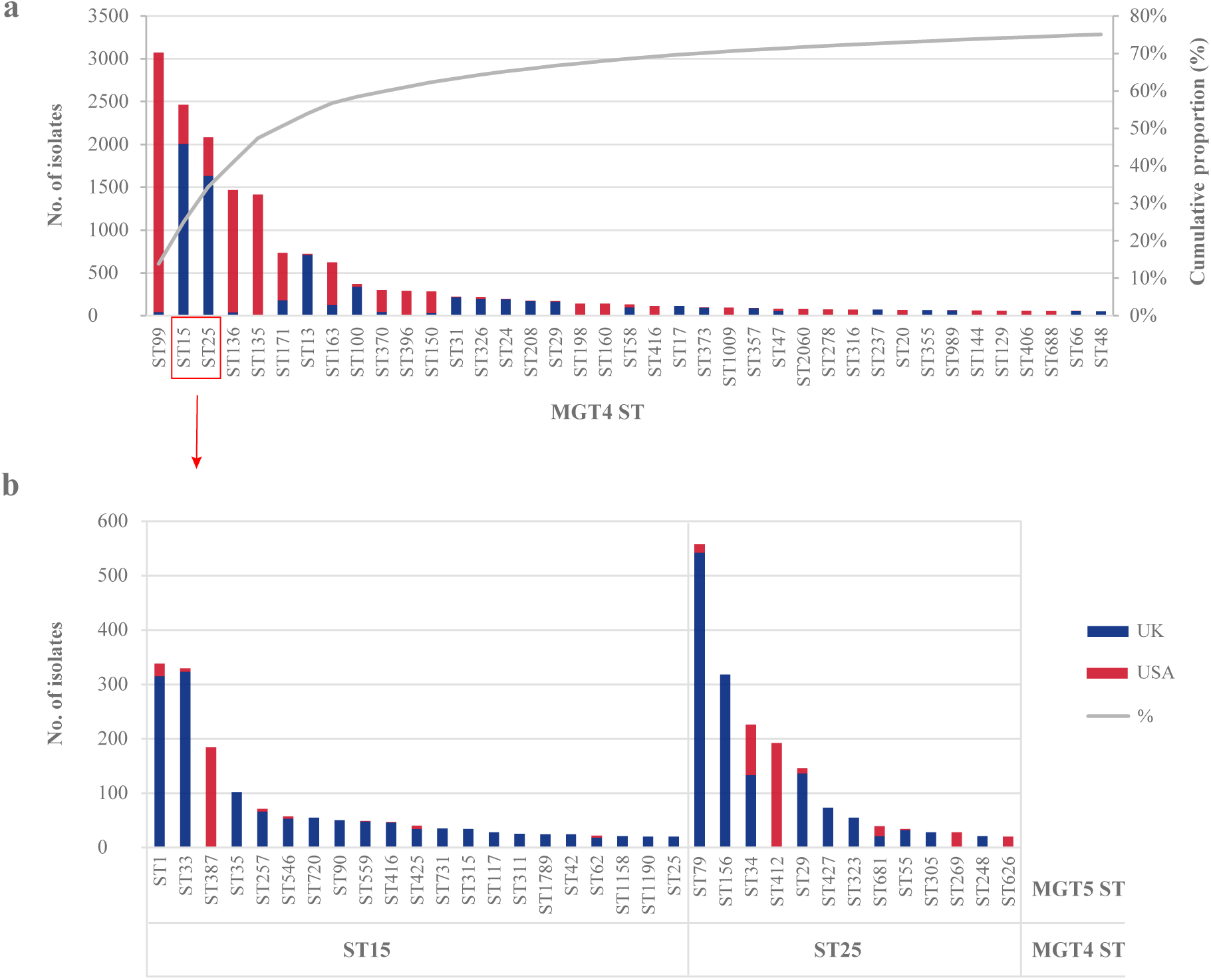
USA and UK *S.* Enteritidis isolates can be distinguished by STs at MGT4 and MGT5. **a.** At MGT4, there were 39 STs (50 or more isolates each) representing 75.1% of the UK and USA isolates (22191 in total). MGT4-ST99, ST136 and ST135 were prevalent in the USA, MGT4-ST15, ST25 and ST13 were prevalent in the UK, while MGT4-ST15 and ST25 were mixed. **b.** MGT4-ST15 and ST25 can be subtyped into multiple STs at MGT5, the majority of which were prevalent in the UK, except for MGT5-ST412 and ST387 in the USA.

To examine the relationship between MGT-STs, isolation source and location, we examined MGT4-STs of 4383 genome sequences from the USA which contained source and state metadata. Of the 4383 genomes, 46.7% were from avian source while 43.1% were from humans. Six MGT4 STs (with more than 50 isolates each) were isolated from both human and avian sources including MGT4-ST99, ST135, ST136, ST160, ST198 and ST25, while three STs including MGT4-ST15, ST163 and ST171 were mainly from human sources (Fig. 4). All of the human-only STs belonged to MGT4-CC1 whereas the mixed source STs were mostly of MGT4-CC13 origin. STs belonging to MGT4-CC13 were significantly more prevalent in avian sources than STs of MGT4-CC1 (P < 0.001, OR = 45.9). For MGT4-CC13, *S.* Enteritidis isolate metadata from 48 states of the USA were available and ST frequencies were similar across the country (Fig. 4). In almost all states, MGT4-ST99 (within MGT4-CC13) was the dominant type, followed by MGT4-ST135 and MGT4-ST136. While for MGT4-CC1 STs, ST15, ST163 and ST171 are predominant in only two states.

**Fig. 4.**
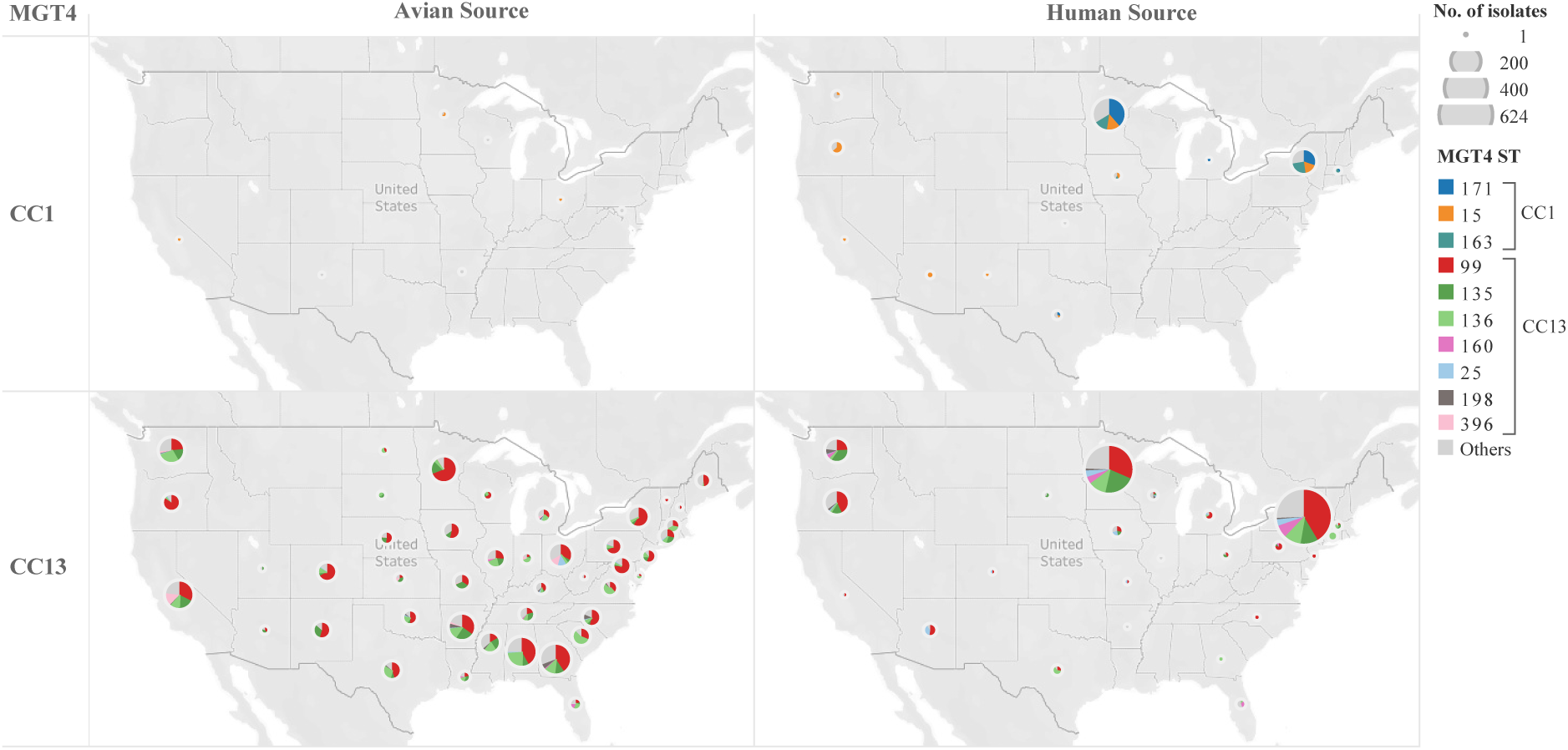
MGT4 ST proportions across states in the USA is similar but only a subset of STs are associated with avian hosts. For the USA *S.* Enteritidis isolates with metadata (4,383 isolates), a total of 3935 genomes were from either avian or human source, 95.0% (3738/3935) of which belonged to MGT4-CC1 and CC13. The MGT4-STs belonged MGT4-CC1 and CC13 of either avian or human source were shown in different states of the USA. Each pie chart illustrated the MGT-ST types and the size of the main STs represented with different colours. The size of the pie indicated the total number of isolates in each state. While CC13 appears to originate in birds before moving to humans, the reservoir of CC1 is unknown. Origins for CC1 STs were also not observed in any other source type. The maps were created with Tableau v2019.2.

Most UK isolates contained collection year and month metadata. There were 8,818 human *S.* Enteritidis isolates collected from March 2014 to July 2018 in the UK. We chose MGT5-STs to describe the monthly variation of *S.* Enteritidis in the UK. By MGT5, variation in the prevalence of MGT5-STs in different years and months was observed (Fig. 5). There were 13 MGT5-STs with more than 100 isolates, representing 46.3% of the 8,818 isolates. The top five STs over the entire period were MGT5-ST1, ST29, ST79, ST15 and MGT5-ST33, representing 30% of the total isolates. These STs showed differing temporal patterns. MGT5-ST1, ST29, ST79 and ST15 were consistently observed in each month across these four years while MGT5-ST15 included isolates previously reported as part of an outbreak [45]. MGT5-ST156 first appeared in April of 2012 in the UK, increased in frequency in June and July of 2014, and became rare after 2014. MGT5-ST423 was a dominant type from March to September of 2015, then became rare and was dominant again from the September of 2016 to January of 2017.

**Fig. 5.**
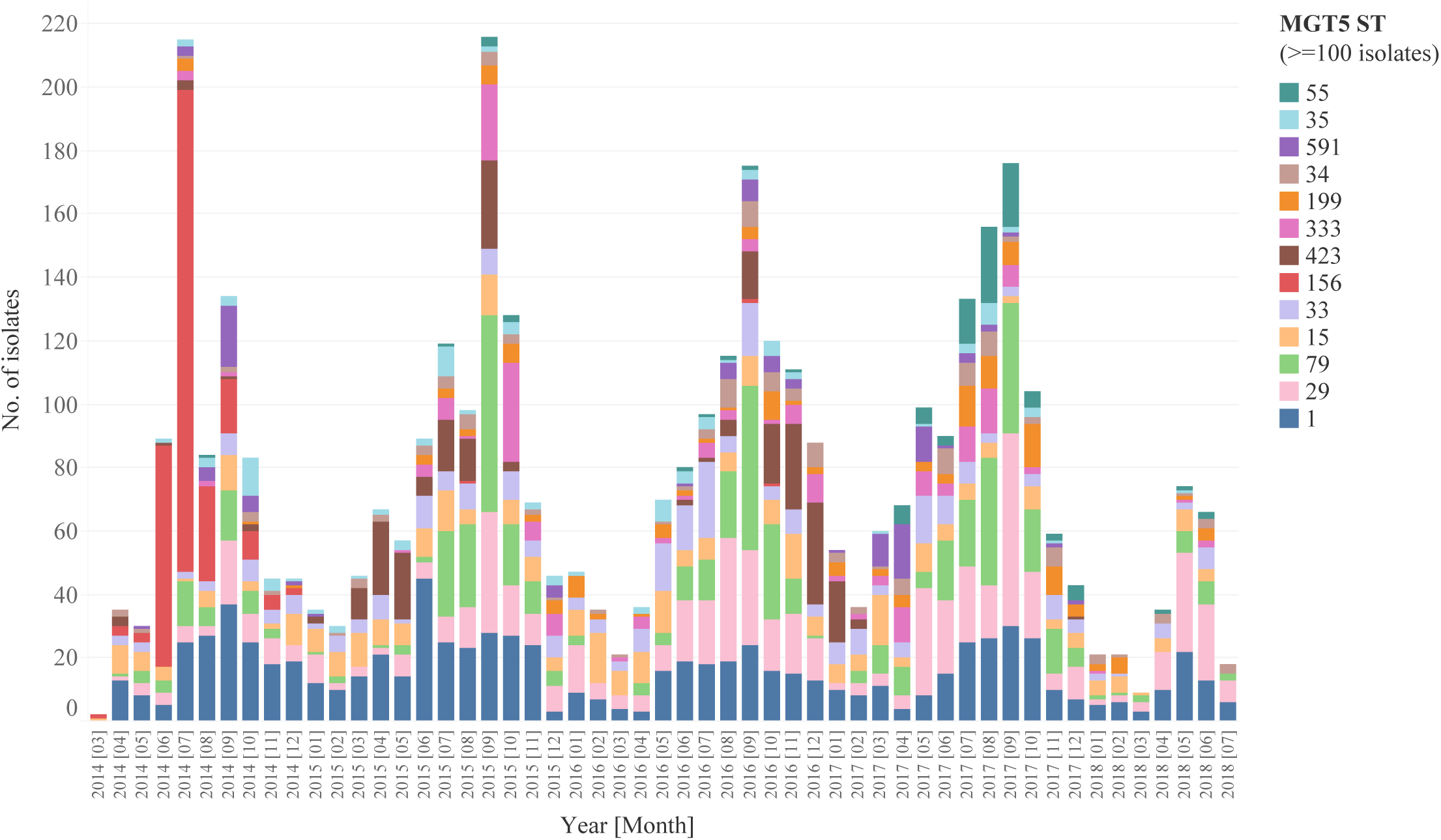
MGT5 STs allow identification of temporal patterns in strain diversity in the UK. There were 13 MGT5-STs of more than 100 isolates shown with different colours. Temporal patterns indicate that some STs are endemic while others likely cause outbreaks. MGT5-ST156 in red was observed predominantly from June to August of 2014, and disappeared after 2014. MGT5-ST423 was a dominant type from March to September of 2015, and became dominant again during the September to December of 2016.

### Detection of potential large outbreak clusters of *S.* Enteritidis using MGT

MGT9 offered highest resolution for tracking strain spread and outbreak detection. By MGT9, there were 124 MGT9-STs including isolates from two or more countries each, indicating international spread (Data Set S1, Tab 3). To facilitate outbreak detection, we identified single linkage clusters of isolates using MGT9 allele differences with a range of cut-offs (0, 1, 2, 5 and 10 allelic differences) that were named as outbreak detection clusters (ODC0, 1, 2, 5 and 10). These clusters can be used to analyse frequencies of closely related isolates at different cut-off levels for population studies and may also be used as dynamic thresholds to detect potential outbreaks [46].

In this study, we used ODC2 clusters to detect potential outbreak clusters and to assess whether some STs were more likely to cause large outbreak clusters, based on the number of ODC2 clusters and the total number of isolates in these clusters in different STs. Since the global data may be biased towards outbreak isolates that were preferentially sequenced, we used UK data from 2014 to 2018 which included all human isolates referred to public health authorities [47]. There were 17 ODC2 clusters of more than 50 isolates representing 1855 isolates in total. The majority of these ODC2 clusters belonged to 12 different MGT4-STs and therefore we used MGT4 level to perform comparison (Data Set S1, Tab 4). MGT4-ST15 (1804 isolates) was the dominant ST in UK and contained only one ODC2 cluster of 62 isolates while MGT4-ST25 (1532 isolates), which was the second dominant ST in UK, contained five large ODC clusters containing 58 to 287 isolates (602 in total). MGT4 ST15 is significantly less likely to contain large ODC2 clusters than the other STs (OR = 0.1, P value < 0.001), while MGT4-ST25 is significantly more likely to contain larger ODC2 clusters than other STs (OR = 3.1, P value < 0.001). It is noteworthy that by clonal complexes, all of the six top STs belonged to MGT4-CC13 and were positively associated with large scale outbreaks, including three previously reported large scale outbreaks in Europe [19, 45, 48, 49]. Indeed, MGT4-CC13 was significantly more likely to cause larger scale outbreaks than MGT4-CC1 (OR = 6.2, P < 0.001). These associations of STs and CCs with large outbreak clusters were also significant when ODC5 clusters was used. Notably the cluster sizes were larger but the trend remains the same (Data Set S1, Tab 4 and Tab 5).

### Antibiotic resistance (AR) gene profiling of *S.* Enteritidis

As nearly all of the isolates (99.98%, 26,665/26,670) harboured the *aac(6’)-Iaa_1* gene for aminoglycoside resistance, this gene was excluded from the antibiotic resistance analysis. A total of 2505 isolates (9.4% of the total 26,670 isolates) were found to harbor antibiotic resistant genes excluding *aac(6’)-Iaa_1*. The most frequent predicted class was tetracyclines (5.1%, 1350/26,670), followed by beta-lactams (4.5%, 1213/26,670) and aminoglycosides (2.8%, 741/2270). Among the 2505 isolates with AR genes, 40.4% (1011/2505) were predicted to be MDR, harbouring genes conferring resistance to three or more different antibiotic classes. And 59.6% (1494/2505) of the isolates harboured genes conferring resistance to one to two different antibiotic classes. Although the selection of isolates for genome sequencing in some continents may be biased, African isolates were found to have the highest proportion of AR/MDR isolates, followed by Asia and Oceania (Fig. S5a).

STs containing more than 10 isolates at different MGT levels were screened to identify MDR or AR associated STs (defined as >= 80% of the isolates are predicted to be MDR or AR). Eleven STs from varied levels were identified as MDR STs, representing 49% (659/1011) of the MDR isolates. These STs were mutually exclusive at different MGT levels. The top two STs, MGT3-ST15 and ST10, were the two invasive types that were prevalent in Africa. For MGT3-ST15, 99.2% of the isolates harboured resistance genes corresponding to as many as six different drug classes (Table 1). For MGT3-ST10, 86.5% of the isolates were MDR, and the antibiotic resistant patterns were similar to those of MGT3-ST15. MGT3-ST10 and ST15 represented 96.3% and 90.2% of the two Africa endemic lineages (MGT3-CC10 and CC15, respectively) as mentioned below. Among the other nine MDR STs, eight belonged to MGT4-CC1, and 93.5% of the isolates (243/260) harboured genes conferring resistance to as many as eight classes of antibiotic drugs. Only one MDR ST (MGT6-ST2698) belonged to MGT4-CC13 with 89.7% (35/39) of the isolates harboured genes conferring resistance to four drug classes. Based on available population sampling, MGT4-CC1 had significantly more isolates with MDR genes (5.6%, 474/8539) than MGT4-CC13 (0.7%, 107/14,972) (Fisher exact test, P value < 0.001, OR = 12.4).

**Table 1.**
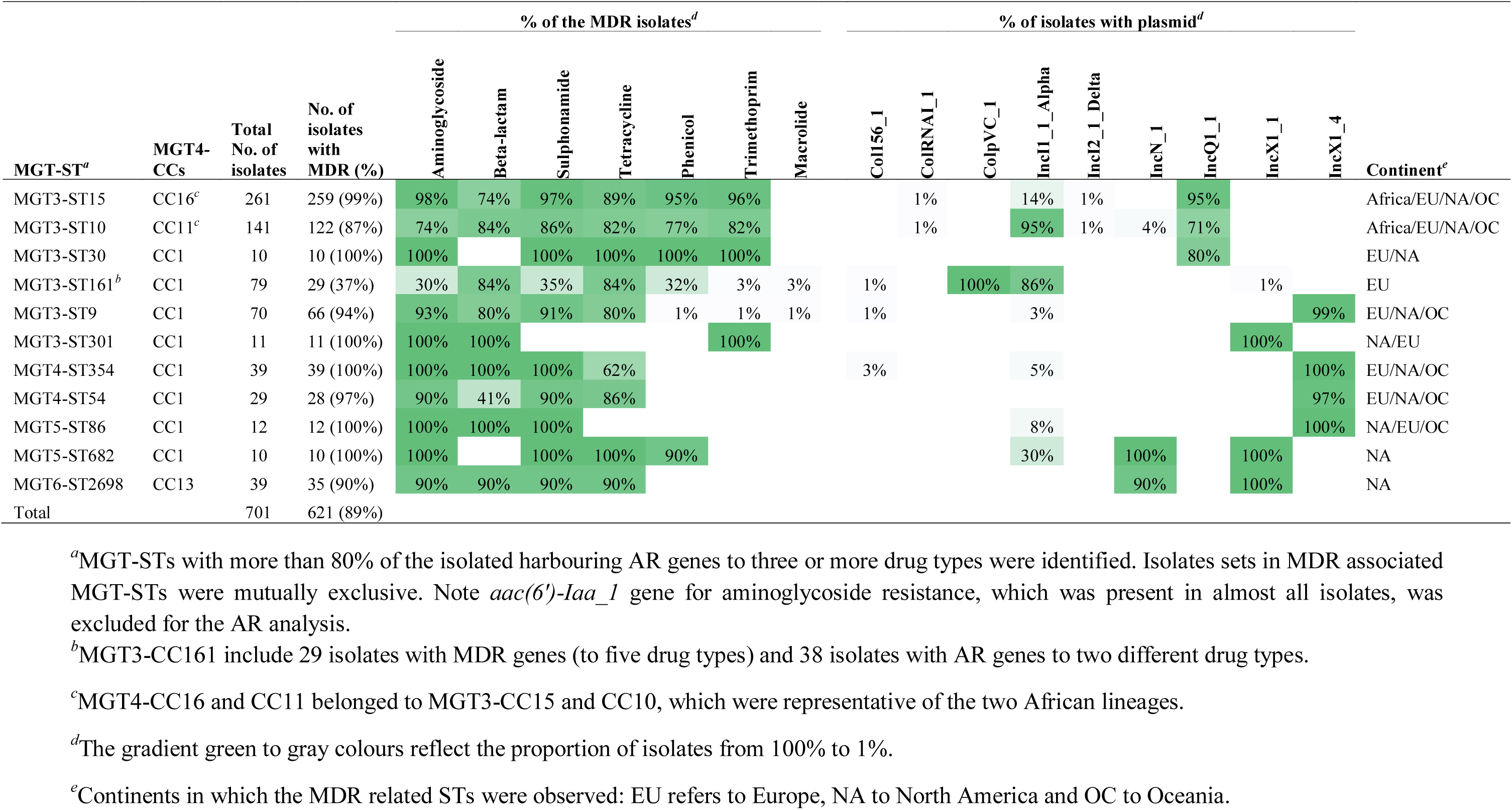
Antibimicrobial drug class and plasmid class of the MDR associated STs from different MGT levels.

A total of 707 isolates belonging to 47 different STs from MGT3 to MGT7, harboured genes conferring resistance to one or two different antibiotic classes including aminoglycosides, beta-lactams, tetracyclines and quinolones (Data Set S1, Tab 6). Among the 47 STs, 34 (72%, 34/47) belonged to MGT4-CC1, representing 80% (564/707) of isolates, and 12 (26%, 12/47) STs belonged to MGT4-CC13 representing 18% (125/707) of the isolates. Again MGT4-CC1 had significantly more isolates with AR genes than MGT4-CC13 (Fisher exact test, P value < 0.001, OR = 8.4).

Plasmid specific genes were identified from the PlasmidFinder database [33] and IncQ1, IncN, IncI1, IncX1 plasmid types were common in *S*. Enteritidis isolates with AR genes (Fig. S5b). In particular, plasmid type IncQ1 was present in MGT3-ST15, ST10 and ST30, which harboured MDR genes up to six drug classes (Table 1). Plasmid type IncI1 was common in MGT3-ST10 and ST161 and plasmid type IncX1 was more common in the STs of MGT4-CC1.

### The population structure and evolution of major *S.* Enteritidis STs/CCs

The majority of the STs from different MGT levels analysed belonged to the two MGT4 CCs, MGT4-CC1 and CC13. To describe the global phylogenetic structure of *S*. Enteritidis and explore the phylogenetic relationship of the two lineages, 1506 representative isolates were selected using representatives of MGT6-STs to encompass the diversity of the serovar. A previous study had suggested that *S*. Enteritidis has three clades, A, B and C. Clade A and C appeared to diverge earlier and were phylogenetically more distant to the global clade B than *Salmonella enterica* serovar Gallinarum [11] (Fig. 6a). *S*. Enteritidis clade B and *S.* Gallinarum were sister clades. The vast majority of the genomes analysed belonged to clade B.

**Fig. 6.**
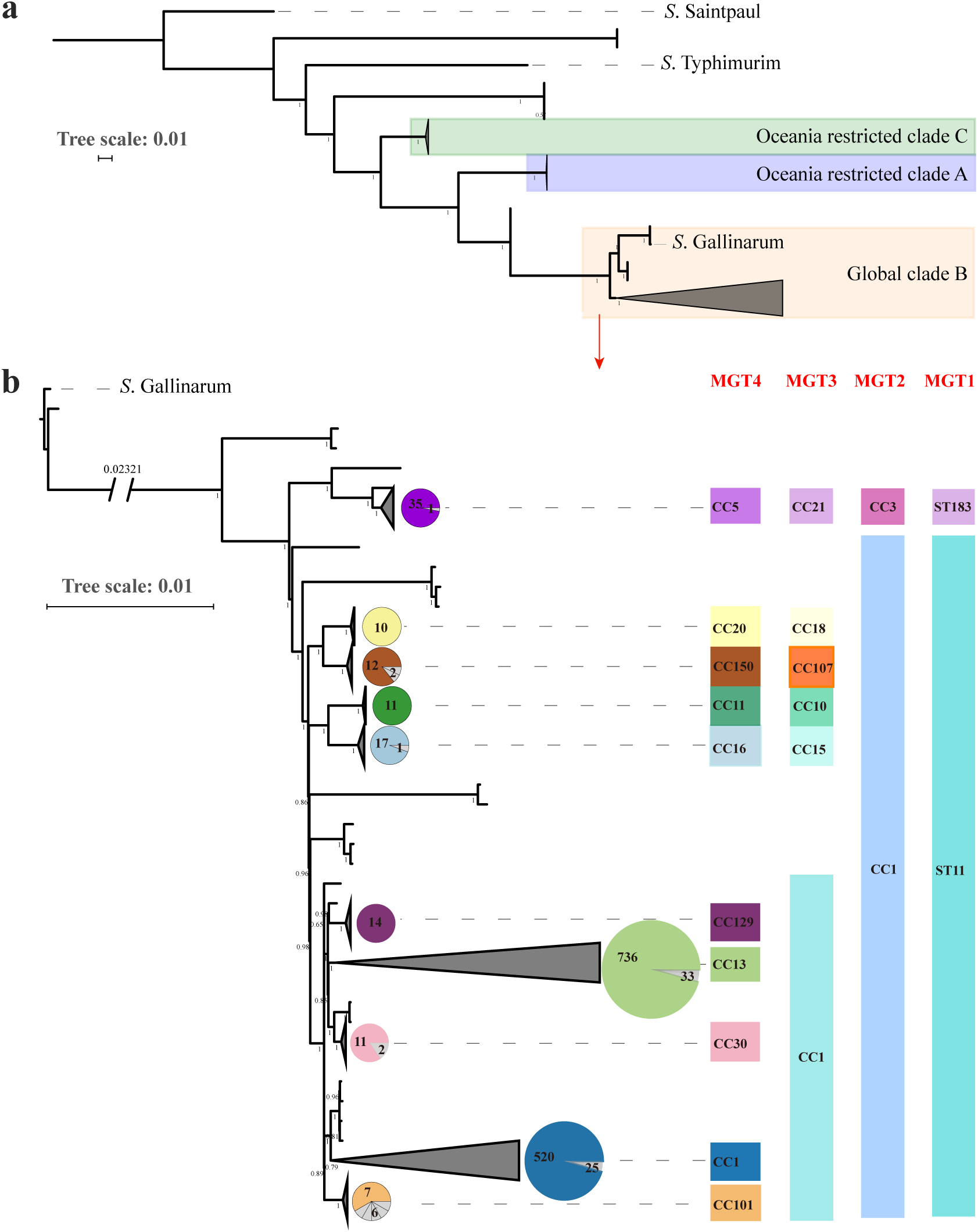
Global phylogenetic structure of *S.* Enteritidis. A total of 1508 genomes were randomly sampled from each ST in MGT6. A phylogenetic tree was constructed by ParSNP v1.2, which called core-genome SNPs extracted from alleles. The tree scales indicated the 0.01 substitutions per locus. The number on internal branches represented the percentage of bootstrap support. **a.** Three clades were identified which were concordant with previous findings: The predominant clade, highlighted in yellow, is the global epidemic clade which was phylogenetically closer to *S.* Gallinarum than the other two clades in purple and green background which are limited in Oceania 9. **b**. The main, yellow highlighted clade was expanded into a more detailed phylogeny with *S.* Gallinarium as an outgroup. 10 main lineages were defined where each lineage had more than 10 isolates. STs of the seven gene MLST (or MGT1) and CCs from MGT2 to MGT4 are shown in different colours for each of the 10 lineages. The pie charts for each lineage represent the proportion of the isolates belonged to the same MGT4-CC shown on the right side.

Within clade B, we identified 10 main lineages, which were concordant with MGT-CCs from MGT1 to MGT4 (Fig. 6b). These lineages can be described at different MGT levels as shown in Fig. 6b. The lineages were consistent with the progressive division from lower to higher MGT levels grouped by CCs. MGT1-ST11 can be represented by MGT2-CC1 and MGT1-ST183 by MGT2-CC3. At MGT3, four lineages named MGT3-CC10, CC15, CC18 and CC107 defined separate lineages, while MGT3-CC1 included several lineages represented by MGT4-CCs. MGT3-CC10 and CC15 represented 100% of the two previously reported Africa lineages associated with invasive infection [5]. MGT3-CC1 included five main MGT4-CCs, with two, MGT4-CC1 and MGT4-CC13, representing 88.0% of all 26,670 *S.* Enteritidis isolates (Fig. 6b). MGT4-CC30 and CC129, both of which were phylogenetically closer to MGT3-CC13, were two endemic lineages in Europe and North America, respectively (Fig. 6b). The population structure of *S.* Enteritidis defined by MGT, were generally in accordance with cgMLST HierCC HC100 (Fig. S6). The association between MGT-STs/CCs with previously reported SNP analysis-based nomenclature (lineages/clades) and phage types were summarized (Dataset 1, Tab 7) [5, 19, 44, 45, 49].

We performed BEAST analysis using the *S.* Enteritidis core genome to determine the evolutionary rate of the clade B *S.* Enteritidis which was estimated to be 1.9×10^-7^ substitution/site/year (95% CI of 1.6×10^-7^ to 2.3×10^-7^), corresponding to 0.8 SNPs per genome per year for the core genome (95% CI of 0.6 to 0.9 SNPs). The mutation rate of MGT4-CC13 lineage core genome was estimated to be 2.5×10^-7^ substitution/site/year (95% CI of 2.3×10^-7^ to 2.7×10^-7^) or 1.0 SNP per genome per year (95% CI of 0.9 to 1.1 SNPs). The mutation rate of MGT4-CC1 lineage core genome was 1.7×10^-7^ substitution/site/year (95% CI of 1.6×10^-7^ to 2.0×10^-7^), or 0.7 SNP per genome per year (95% CI of 0.6 to 0.8 SNPs). The mutation rate of MGT4-CC13 was significantly faster (1.5 times) than that of MGT4-CC1.

The most recent common ancestor (MRCA) of the nine lineages belonging to MGT2-CC1, was estimated to have existed in the 1460s (95% CI 1323 to 1580) (Fig. S7). The two global epidemic lineages MGT4-CC1 and CC13 are estimated to have diverged at around 1687 (95% CI 1608 to 1760). In 1869 (95% CI=1829-1900), MGT4-CC13 diverged into two sub-lineages of various MGT4-STs, with one more prevalent in North America than in other continents (labelled with red arrow) and the other sub-lineage more prevalent in Europe (labelled with blue arrow). We further estimated the population expansion of the two global epidemic lineages MGT4-CC1 and CC13 (Fig. 7a, b). For MGT4-CC1, there were two large expansions around 1950 and 1970. For MGT4-CC13, the population gradually increased until a more rapid expansion around 1970.

**Fig. 7.**
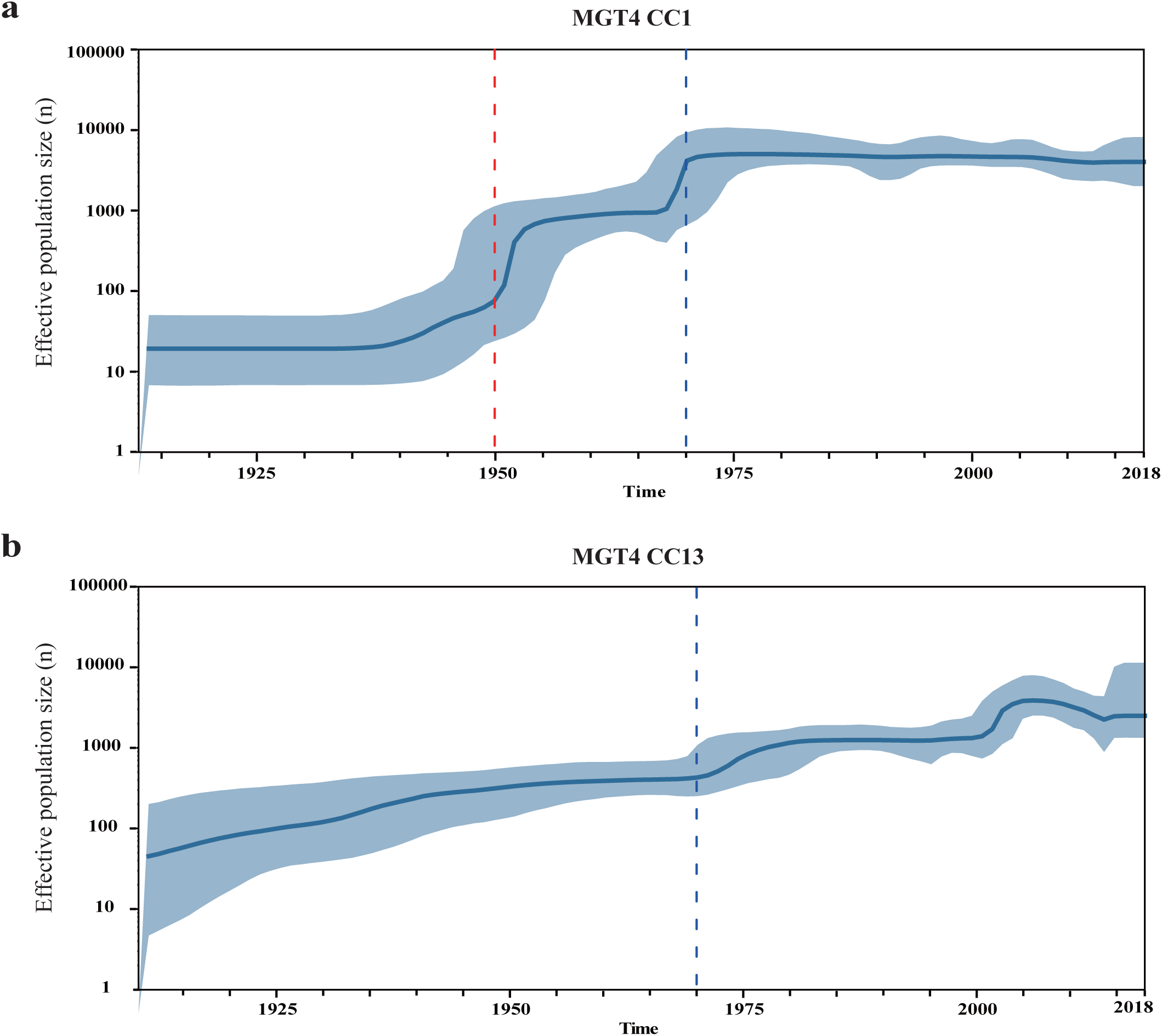
Skyline population size estimation of MGT4-CC1 and CC13. Bayesian skyline model and strict clock were estimated to be the optimal combination. The vertical axis column indicates the predicted log scale effective population size and the horizontal axis shows the predicted time. The blue line represents the median posterior estimate of the effective population size, and the blue area shows the upper and lower bounds of the 95% highest posterior density interval. **a.** Skyline population expansion of MGT4-CC1. Around 1950 and 1970, there were two large population expansions within MGT4-CC1. **b.** Skyline population expansion of MGT4-CC13. The population of MGT4-CC13 gradually expanding until around 1970, when there was also an accelerated expansion.

### Virulence genes and SPIs distribution in the STs/CCs represented phylogeny of *S.* Enteritidis

We further compared the distribution of virulence genes and SPIs in the 13 STs/CCs that represent the major phylogenetic lineages of *S.* Enteritidis. Based on the VFDB database, 162 genes were present in >= 10 isolates of *S.* Enteritidis, 123 (75.9%) genes of which were present in all of the STs/CCs. Sixteen genes (9.9%) were associated with one or more STs/CCs while the remainder were sporadically distributed (Table 2). The *spv* locus including *spvB*, *spvC* and *spvR*, are reported to be associated with non-typhoidal bacteraemia [50]. *SpvBCR* genes were absent in MGT1-ST3304, ST180, ST1972 but present in all the other CCs. Pef fimbriae operon *pefABCD* genes were absent in MGT1-ST3304, ST180, ST1972, ST183 and MGT3-CC10. The *ssek2* gene, which encodes a secretion effector of SPI-2, was reported to significantly contribute to the pathogenicity of *Salmonella* [51]. *ssek2* was present in MGT3-CC107 and CC18, MGT4-CC129 and CC1, but was absent in MGT4-CC13 and other STs/CCs. On the reference genome P125109 (MGT4-CC1), *ssek2* was observed in the prophage □SE20. Moreover, 34% of the isolates in MGT3-CC107 (further represented by five STs from MGT4 to MGT7 levels) were found to harbour the *Yersinia* high-pathogenicity island (HPI), representing 75.5% of the HPI positive isolates (Data Set S1, Tab 8).

**Table 2.**
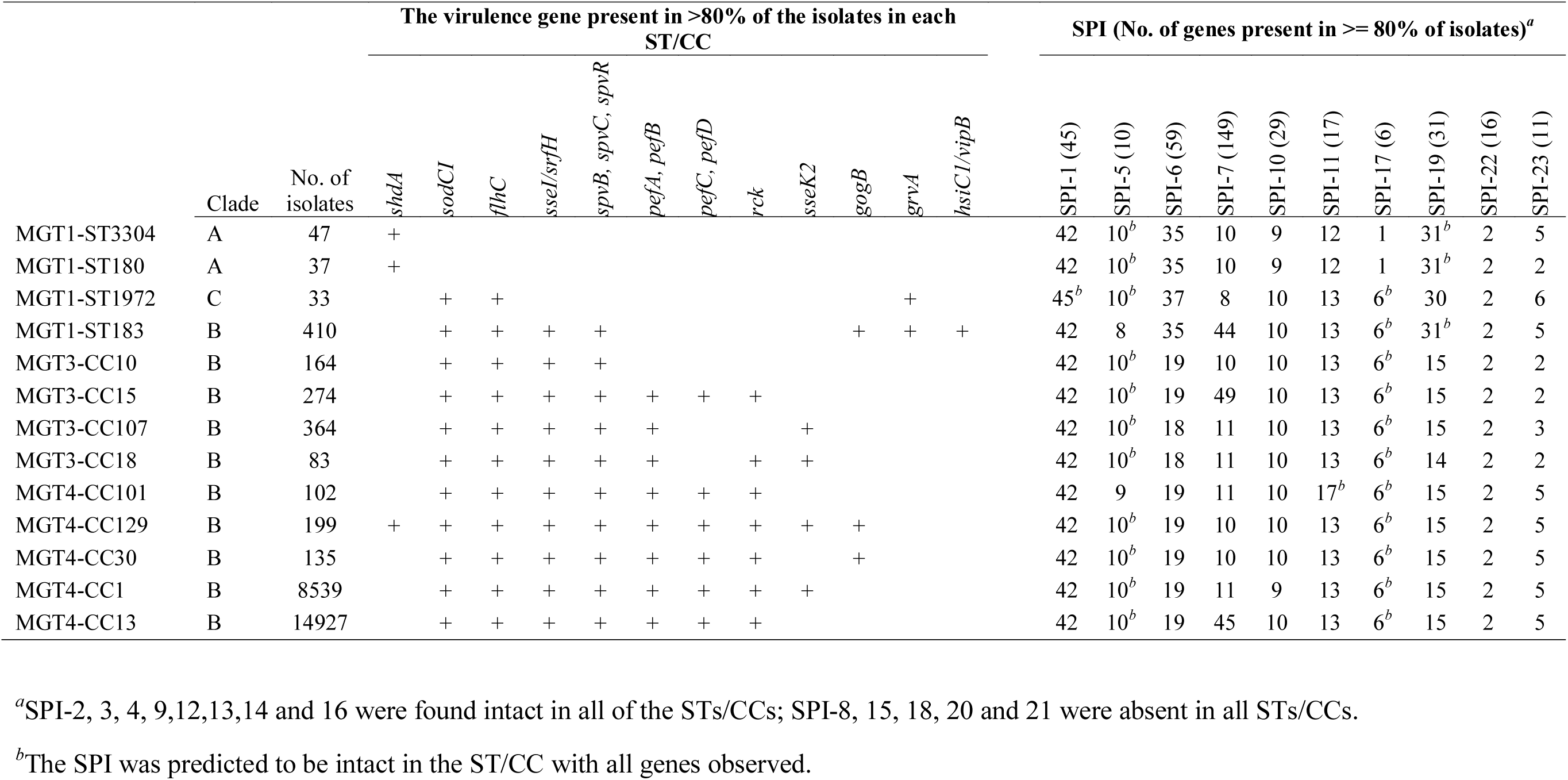
Virulence and SPI genes that distributed differently in the STs/CCs represented phylogeny of *S.* Enteritidis.

Among the 23 SPIs reported, SPI-2, 3, 4, 9,12,13,14 and 16 were found intact in all of the STs/CCs, and SPI-8, 15, 18, 20 and 21 were absent in all STs/CCs (Table 2). SPI-1 was intact in MGT1-ST1972 (clade C), while all the other STs/CCs had three SPI-1 genes (STM2901, STM2902 and STM2903) missing. SPI-5 was nearly intact in all STs/CCs with one to two genes missing in MGT1-ST183 and MGT4-CC101. SPI-11 was only intact in MGT4-CC101, with four to five genes missing in the other STs/CC. SPI-17 was intact in the majority of the STs/CCs except for MGT1-ST3304 and ST180 (clade A) with only one gene observed. SPI-19 was nearly intact in MGT1-ST3304, ST180, ST1972 and ST183, but was truncated in all the other CCs in clade B. None of the STs/CCs harboured the intact SPI-6, 7, 10, 22, and 23, only a few genes of which were observed. MGT1-ST183, MGT3-CC15 and MGT4-CC13 were found to harbour 44 to 49 genes of SPI-7 (149 in total), the majority of which were located on the Fels2-like prophage.

## Discussion

*S.* Enteritidis is one of the most common foodborne pathogens causing large scale national and international outbreaks and food recalls [3, 52]. Understanding the population structure and genomic epidemiology of *S.* Enteritidis is essential for its effective control and prevention. In this study, we applied the genomic typing tool MGT to *S.* Enteritidis and developed a database for international applications. We used MGT to describe its local and global epidemiology, and its population structure using 26,670 publicly available genomes and associated metadata. In this work, STs and CCs were assigned to each of the nine MGT levels, with MGT1 refers to the legacy seven gene MLST [12]. Thus, attention should be paid to the prefix MGT levels for those STs and CCs to avoid confusion.

### The *S.* Enteritidis MGT scheme enables a scalable resolution genomic nomenclature

The design of the *S.* Enteritidis MGT is based on the *S.* Typhimurium MGT scheme published previously [20]. The first eight levels (MGT1 to MGT8) of *S.* Enteritidis MGT scheme used the same loci as for *S.* Typhimurium. We defined the core genome of *S.* Enteritidis which had 3932 core genes and 1054 core intergenic regions, which was used as MGT9. MGT9 substantially increased the subtyping resolution for *S.* Enteritidis [13]. Our study suggests that the MGT levels one to eight could be applied to all *Salmonella enterica* serovars as a common scheme for the species, and only an additional serovar-specific MGT9 scheme needs to be designed for individual serovars that require the highest resolution for outbreak investigations.

The online database of the *S.* Enteritidis MGT scheme offers an open platform for global communication of genomic data and facilitates detection of international transmission and outbreaks of *S.* Enteritidis. The STs, especially at the middle resolution MGT level, were associated with different geographic regions, sources and MDR. The extension of MGT to *S.* Enteritidis was based on our previous study on *S.* Typhimurium [20]. Importantly, the stable characteristics of STs, which are based exact comparison, makes up for the main drawback of single-linkage clustering methods that the cluster types may change, especially at higher resolutions. MGT STs enables long term epidemiological communication between different laboratories. And the variable resolution of MGT offers flexibility for temporal and spatial epidemiological analysis using an underlying stable nomenclature of STs from the finest resolution level. Additionally, CCs at each MGT level were able to cluster the STs with one allele difference and are concordant with phylogenetic lineages as shown in this study.

### MGT for *S.* Enteritidis uncovers geographic, source and temporal epidemiological trends of *S.* Enteritidis within and between countries

A total of 26,670 isolates were successfully typed using the MGT scheme which allowed the examination of the global (or international) and local (or national) epidemiology of *S.* Enteritidis. The salient feature of the flexibility of the different levels of MGT has also been illustrated through this analysis. The lower resolution levels of MGT were found to be able to effectively describe the geographic variation in different continents, countries or regions. Of these levels we showed that MGT4-STs best described the global epidemiology of *S.* Enteritidis at continental level with some STs distributed globally while others more geographically restricted. MGT4-ST15 was a global epidemic type which was prevalent in almost all continents. By contrast, MGT4-ST16 (also described by MGT3-ST15) was prevalent in Central/Eastern Africa and MGT4-ST11 (also described by MGT3-ST10) was prevalent in West Africa, which agreed with a previous study [5].

In USA, MGT4-CC13 and MGT-CC1 showed remarkable difference in their epidemiology. MGT4-CC13 STs including MGT4-ST99, ST136 and ST135 were the main cause of clinical infections and were commonly isolated from poultry. The prevalent STs were generally similar in different states. In particular, MGT4-ST99, ST136 and ST135 were the dominant types in almost all states of the USA. The distribution and poultry association of these STs suggest that they may be responsible for several multistate outbreaks caused by *S*. Enteritidis contaminated eggs [1, 53]. Since poultry related products (i.e. eggs, chicken and turkey, especially eggs) are known to be the main source of *S.* Enteritidis infections or outbreaks in the USA [4], this isn’t surprising, but it demonstrates the utility of the MGT.

On the other hand, MGT4-CC1 including MGT4-ST171, ST15 and ST163, were less prevalent in the USA. Nevertheless, these STs were isolated from human infections but were very rare in poultry, although sampling bias may affect this conclusion as the poultry isolates in the dataset was not from systematic sampling of poultry sources. On the other hand, beef, sprouts, pork, nuts and seeds can also be contaminated by *S.* Enteritidis [4]. These non-poultry related foods may be the source of the MGT4-CC1 caused infections. Overall MGT4-CC13 was found to be significantly associated with poultry source, which is concordant with a previous study [10]. Further studies are required to explain the association of MGT4-CC13 but not MGT4-CC1 with poultry products. The lack of potential source isolates within MGT4-CC1 STs highlights the need for comprehensive sequencing and epidemiological efforts across the food production chain. Machine learning approaches, which have been applied to the root source identification for *S.* Typhimurium outbreaks, could also facilitate the source tracing of *S.* Enteritidis [17].

Temporal variation of UK *S.* Enteritidis was depicted clearly by MGT. From 2014 onwards, all clinical *S.* Enteritidis were routinely sequenced in the UK [47]. Some MGT5-STs were found to occur for a few months and then disappear and may be indicative of outbreaks. For example, MGT5-ST156, which increased substantially during June and July in 2014 but became rare after 2015 and describes a reported large scale outbreak [19]. Other STs persisted for long periods of time. For example, MGT5-ST1 and ST29, appeared to be endemic isolates which may be associated with local reservoirs. Therefore, the variation of MGT-STs across different seasons offered additional epidemiological signals for suspected outbreaks and endemic infections of *S.* Enteritidis. The stability of STs avoids the potential cluster merging issues of single linkage clustering based methods and ensures continuity for long-term surveillance [20].

### MGT for *S.* Enteritidis facilitates outbreak detection, source tracing and evaluation of outbreak propensity

Whole genome sequencing offered the highest resolution for outbreak detection and source tracing. However, resolution of cgMLST heavily depends on the diversity of the species. Our previous study showed that for *S.* Typhimurium, serovar core genome offered higher resolution than species level core genome for outbreak investigation [20, 54]. We designed MGT9 for *S.* Enteritidis using 4986 loci, which is around 2000 more loci than *Salmonella enterica* core genes [13, 20], thus increasing the resolution of subtyping.

To facilitate outbreak detection MGT9 STs were further grouped to ODCs. There is no agreed upon single cut-off for outbreak detection and dynamic thresholds have been suggested [46]. Here, we used ODC2, which has a two-allele difference cut-off, to identify potential outbreak clusters. Although the data do not allow us to confirm whether any of these ODC2 clusters were actual outbreaks, a number of known outbreaks from other studies fell into ODC2 clusters [46, 48, 55]. A two-allele cut-off was selected because it is at the lower end of cut-offs in reported Enteritidis outbreaks [48, 55], which should limit the number of false positive outbreak calls. However, due to the variability in diversity of isolates from different outbreaks, a dynamic threshold for cut-offs would be more sensitive and specific to detect outbreaks [46]. Further work for *S.* Enteritidis is required to address this issue fully. Using the UK clinical isolates from 2014 to 2018, we found that the European and North American prevalent lineage MGT4-CC13 was significantly correlated with larger ODC2 clusters (>= 50 isolates each) than the global epidemic lineage of MGT4-CC1. Thus, MGT4-CC13 is more likely to cause large scale outbreaks than MGT4-CC1. In the past few years, several large scale outbreaks in Europe were due to the contaminated eggs [48, 49, 52]; these isolates all belonged to MGT4-CC13. In the USA, MGT4-CC13 was the dominant lineage in both human infections and poultry product contamination, while MGT4-CC1 was relatively rare in poultry. Industrialised and consolidated poultry/eggs production and marketing could have facilitated the spread of MGT4-CC13 causing large scale outbreaks. Further studies are required to definitively identify the biological and environmental mechanisms facilitating these large MGT4-CC13 outbreaks.

ODCs offer a means for accurate detection of outbreaks at the highest typing resolution, MGT9 [20]. Other studies used SNP address or cgMLST [19, 45, 48, 49] with SNP cut-offs or HierCC clustering for outbreak detection. All three methods use the same single linkage clustering method for outbreak detection. MGT offers further advantage that potential outbreak clusters can be precisely defined by a stable MGT-ST identifier rather than a cluster number [20].

### MGT for *S.* Enteritidis improves precision of detection and tracking of MDR clones globally

The rise of AMR in *Salmonella* is a serious public health concern [5]. The global spread of AMR can be mediated by lateral transfer of resistance genes as well as clonal spread of resistant strains. This study systematically evaluated the presence of antibiotic resistant genes in the 26,670 genomes of *S.* Enteritidis, 9.4% of which harboured antibiotic resistant genes. Among the two global epidemic lineages, MGT4-CC1 was found to contain significantly more antibiotic resistant isolates than MGT4-CC13.

We further identified STs that were associated with MDR. Eleven STs (>= 10 isolates each, 659 isolates in total) of different MGT levels were associated with MDR. These STs from different MGT levels were mutually exclusive, emphasising the flexibility of MGT for precise identification of MDR clones. MGT3-ST15 and MGT3-ST10 were representative of the previously reported Africa invasive infection related lineages [5], harboured resistant genes of up to six different antibiotic classes. The other STs, which were mainly from Europe and North America, harbour AR genes to up to eight different classes of drugs. MGT3-ST30 and MGT4-ST718, which belonged to MGT4-CC1 and were observed in Europe, had been reported to be MDR phenotypically [56]. Significantly, isolates from these two STs were mainly isolated from blood samples [56]. These MDR STs correlated with blood stream infections should be monitored closely as they can potentially be more invasive [5].

Because plasmids are known to play a key role in the acquisition of drug resistance genes [57, 58], we identified Inc plasmid groups that are likely present in the *S.* Enteritidis population. Several of these putative plasmids were MDR associated. MDR of African isolates in MGT3-ST10 and ST15 has been reported to be mediated by plasmids [5]. The West Africa lineage MGT3-ST10 isolates were reported to have IncI1 plasmids [59]. In this study, both IncI1 and IncQ1 plasmid were present in the Africa MGT3-ST10 and ST15 isolates as well as STs belonged to global epidemic lineage MGT4-CC1. IncQ1 plasmids, which are highly mobile and widely transferred among different genera of bacteria [60], are correlated with MDR of *Salmonella* [61, 62]. IncI1 plasmids are responsible for the cephalosporin resistance of *Salmonella* and *Escherichia* in several countries [63–65]. Moreover, IncN and IncX1 plasmids were common in the MDR associated STs of MGT4-CC1 and CC13. The use of MGT could enhance the surveillance of drug resistance plasmids.

### Defining the population structure of *S*. Enteritidis with increasing resolution and precision at different MGT levels

In this study, we defined the global population structure of *S*. Enteritidis using MGT-CCs. Three clades of *S*. Enteritidis were previously defined with clade A and C as outgroups to the predominant clade B [11], a classification supported by this study. We further identified 10 lineages within clade B, which can be represented by different CCs from MGT2 to MGT4 corresponding with time of divergence. Lower resolution level MGT CCs described old lineages well while higher resolution level MGT CCs described more recently derived lineages (Fig. S7). Thus, the epidemiological characteristics of a lineage can be identified using STs and/or CCs at appropriate MGT level. MGT2-CC3 represent the Europe endemic MGT1-ST183 lineage, within which MGT2-ST3 and MGT2-ST82 were able to identify previously defined phage types PT11 and PT66 respectively [44]. MGT3-CC10 included all of the reported West Africa lineage isolates, and MGT3-CC15 included all of the reported Central/Eastern Africa lineage isolates as well as Kenyan invasive *S*. Enteritidis isolates [5, 8]. MGT3-ST10 and MGT3-ST15 (represented 96.3% of MGT3-CC10 and 90.2% of MGT3-CC15 respectively), were MDR with similar resistance patterns. Most importantly, the two major global epidemic lineages are clearly described by MGT4-CC1 and MGT4-CC13. MGT4-CC1 corresponds to the global epidemic clade in the study of Feasey *et al*., and MGT4-CC13 to the outlier clade of that study [5]. MGT4-CC1 and CC13 correspond to lineage III and lineage V in Deng *et al*’s study [10]. Thus, by using STs and/or CCs of different MGT levels the population structure of *S.* Enteritidis can be described clearly and consistently.

Interestingly, the two dominant lineages, MGT4-CC1 and CC13, showed different population expansion time and evolutionary dynamics. Bayesian evolutionary analysis revealed a core genome mutation rate for *S.* Enteritidis of 1.9×10^-7^ substitution/site/year which is similar to the genome mutation rate of 2.2×10^-7^ substitution/site/year estimated by Deng *et al* [10]. In contrast, the mutation rate of MGT4-CC13 was 1.5 times faster than MGT4-CC1. Population expansion of MGT4-CC1 occurred with two peaks in the 1950s and 1970s, while MGT4-CC13 population increased steadily until the 1970s. The expansion around the 1970s may have been driven by the development of the modern industrialised poultry industry, including poultry farms and/or processing plants, that *S.* Enteritidis colonized. This is concordant with Deng *et al*’s speculation that the expansion of *S.* Enteritidis population was correlated with the poultry farm industry [10, 18].

### Acquisition and degradation of virulence factors in the STs/CCs represent phylogeny of *S.* Enteritidis

By screening the virulence genes and SPIs in different STs/CCs representing the three clades and major lineages of *S.* Enteritidis, distributional differences were observed for some virulence genes and SPIs. The main difference observed was between clade A/C and clade B, and MGT1-ST183 and other lineages of clade B. *SpvBCR*, *pefABCD*, and *rck* genes were found to be located on the same plasmid in the complete genomes of *S.* Enteritidis (data not shown). However, the Europe endemic MGT1-ST183 and the West Africa endemic MGT3-CC10 were positive for the *spvBCR* genes, but were negative for the *pefABCD* and *spvBCR* genes, suggesting that there may exist other mechanisms of acquisition of *spvBCR* genes. Variation in the distribution of genes on SPIs was also observed. Some virulence or SPIs genes were likely to have been lost in certain STs/CCs. Degradation of virulence genes had been previously observed in *S.* Enteritidis and *S.* Typhimurium [5, 66]. In summary, both acquisition and degradation of virulence factors occurred in the evolution of *S*. Enteritidis.

Moreover, the main differences in virulence factors between the two dominant lineages MGT4-CC1 and CC13 were *ssek2* and the number of SPI-7 genes. *ssek2*, located on the prophage □SE20 was limited to MGT4-CC1 [5], and around 35 SPI-7 genes, located on the Fels2-like prophage were limited to MGT4-CC13 [5]. □SE20 was found to contribute to mouse infections [67]. Fels2-like prophage was also present in a bloodstream infection related lineage three of *S.* Typhimurium ST313 in Africa, which is estimated to have significantly higher invasiveness index than the other lineages of ST313 [66]. However, the detailed pathogenic role of Fels2-like prophage in *Salmonella* infections remained unclear. There are potentially undiscovered virulence factors contributing to the epidemiological variation (infection severity, outbreak propensity and geographic spreading) among clades and lineages of *S.* Enteritidis.

### Limitations of this study and challenges for public database

The epidemiological results generated in this study were based on the publicly submitted metadata and the correctness of the results therefore depends on the accuracy of this data. In as many cases as possible metadata were verified from published sources. However, the possibility exists that incorrect metadata or repeated sequencing of the same isolates influence the results. We showed the effect of the latter was minimal (**Supplementary Material**). However, good metadata is essential, and an internationally agreed on minimal metadata would be useful for better epidemiological surveillance of *S.* Enteritidis. The online MGT system offers unprecedented power for monitoring *S.* Enteritidis across the globe. Good metadata further enhances that power. Moreover, all the genomes of this work were Illumina short-read sequencing except for one reference complete genome. The thresholds of identity and coverage for searching AR genes and plasmid replicon genes were >= 90%. Incomplete genomes and these thresholds may result in some AR and plasmid replicon genes being missed. Additionally, considering the mobile nature of plasmids and prophages, some STs may lose AR or virulence genes in rare cases, timely updating of ST definitions with respect to AR and virulence states will therefore be necessary.

## Conclusions

In this study, we defined the core genome of *S.* Enteritidis and developed an MGT scheme with nine levels. MGT9 offers the highest resolution and is suitable for outbreak detection whereas the lower levels (MGT1 to MGT8) are suitable for progressively longer epidemiological timescales. At the MGT4 level, globally prevalent and regionally restricted STs were identified, which facilitates the identification of the potential geographic origin of *S.* Enteritidis isolates. Specific source associated STs were identified, such as poultry associated MGT4-STs, which were common in human cases in the USA. At the MGT5 level, temporal variation of STs was observed in *S.* Enteritidis infections from the UK, which reveal both long lived endemic STs and the rapid expansion of new STs. Using MGT3 to MGT6, we identified MDR-associated STs to facilitate tracking of MDR spread. Additionally, certain plasmid types were found to be highly associated with the same MDR STs, suggesting plasmid-based resistance acquisition. Furthermore, we evaluated the phylogenetic relationship of the STs/CCs by defining the population structure of *S*. Enteritidis. A total of 10 main lineages were defined in the globally distributed clade B of *S*. Enteritidis, which were represented with 10 STs/CCs. Of these, MGT4-CC1 and CC13 were the dominant lineages, with significant differences in large outbreak frequency, antimicrobial resistance profiles and mutation rates. Some virulence and SPIs genes were distributed differently among the different STs/CCs represented clades/lineages of *S.* Enteritidis. The online open database for *S.* Enteritidis MGT offers a unified platform for the international surveillance of *S.* Enteritidis. In summary, MGT for *S*. Enteritidis provides a flexible, high resolution and stable genome typing tool for long term and short term surveillance of *S*. Enteritidis infections.

## Authors’ contributions

L.L., M.P., and R.L. designed the study. L.L. and M.P. set up the database. S.K and M.P. set up the website, M.P., R.L., D.H. L.C. S.O., Q.W., M.T., V.S. and S.K. provided critical analysis and discussions, L.L. wrote the first draft and all authors contributed to the final manuscript.

## Supporting information

Supplementary Figures and Results

Data Set S1

Data Set S2

## Acknowledgements

The authors thank Duncan Smith and Robin Heron from UNSW Research Technology Services for high performance computing assistance.

## Abbreviations

MGT: Multilevel genome typing;
ST: Sequence type
CC: Clonal complex
ODC: Outbreak detection cluster
SNP: Single nucleotide polymorphism
MLST: Multi-locus sequence typing
cgMLST: core genome multi-locus sequence typing
HierCC: Hierarchical clustering of cgMLST
AR: Antibiotic resistance
MDR: Multi-drug resistance

