## Supplementary Figures and Results for "Elucidation of global and local genomic epidemiology of *Salmonella enterica* serovar Enteritidis through multilevel genome typing"

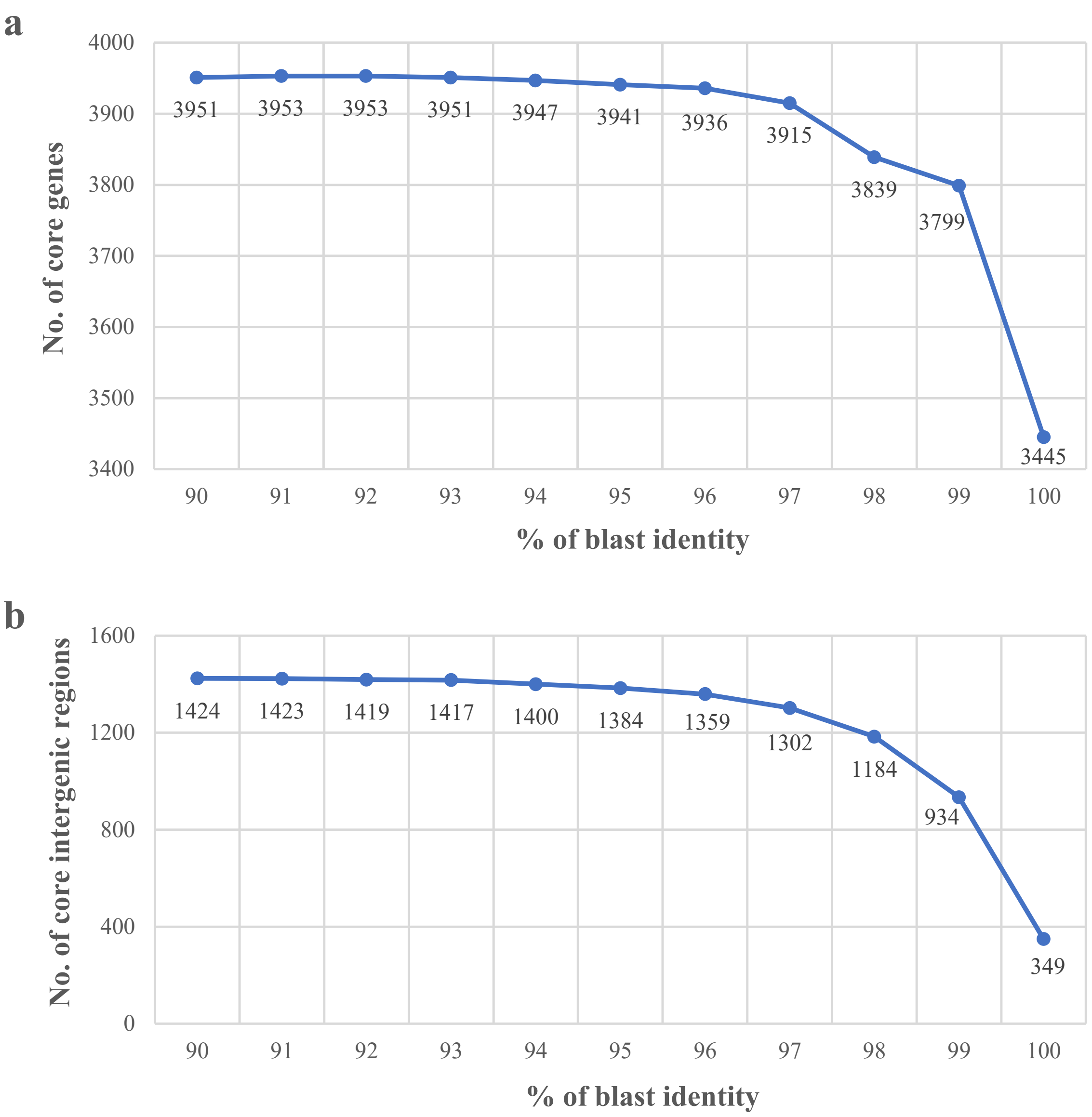


**Fig. S1. Blast identity and the number of core genes and core intergenic regions defined for *S.* Enteritidis. a**. Number of core genes defined by Roary v3.11.2 with varied blast identity from 90% to 100% and gene coverage of 99%. **b**. Number of core intergenic regions defined by Piggy v3.11.2 with varied blast identity from 90% to 100% and coverage of 99% using the Roary output.

**a**

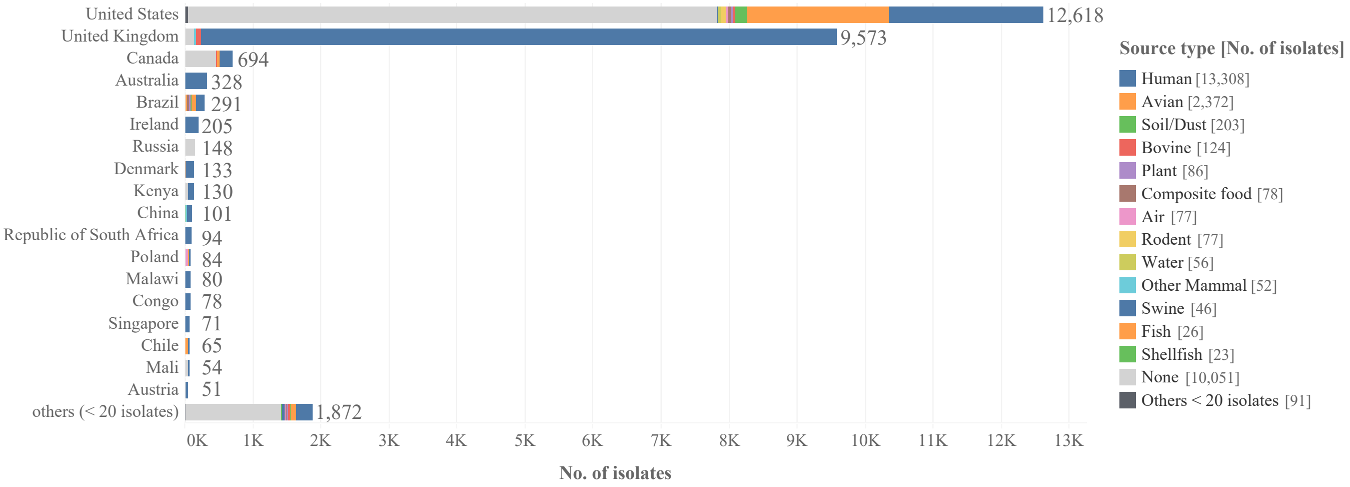


**b**

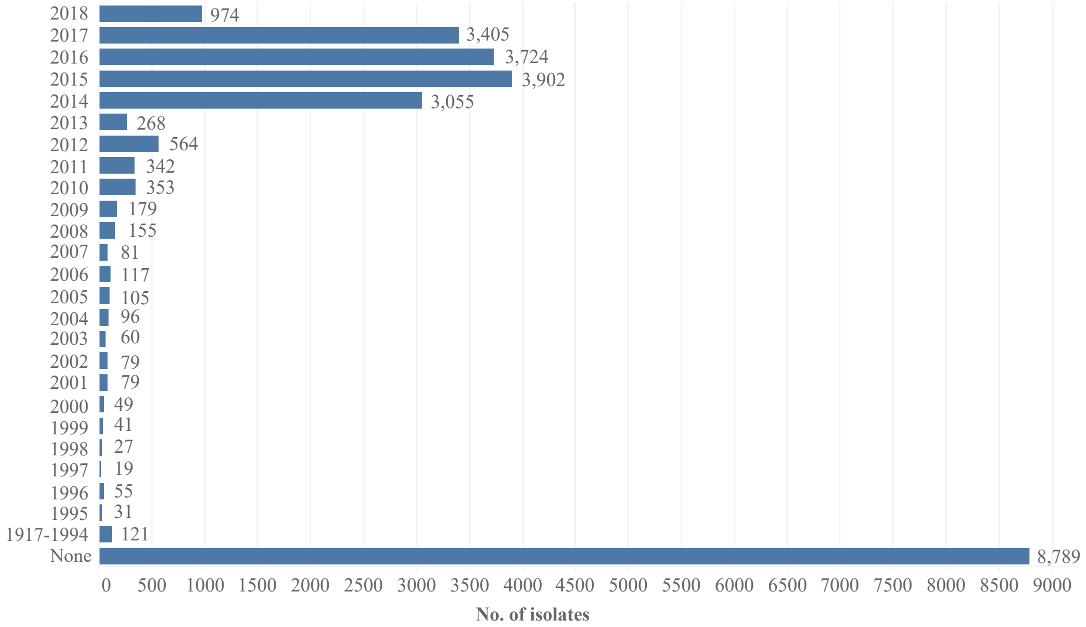


**Fig. S2. General epidemiological information for the 26670 isolates analysed with the following general metadata information**: **a**. isolated from 86 different countries [with the majority of the isolates isolated from the United States (47.3%) and United Kingdom (35.9%)]; isolated from 26 different source types (the source types of each country were shown in different colours); **b**. collected between 1917 and 2018 [with 2014 (11.5%), 2015 (14.6%), 2016 (14.0%), 2017 (12.8%) and 2018 (3.7%) having more than 500 isolates each].

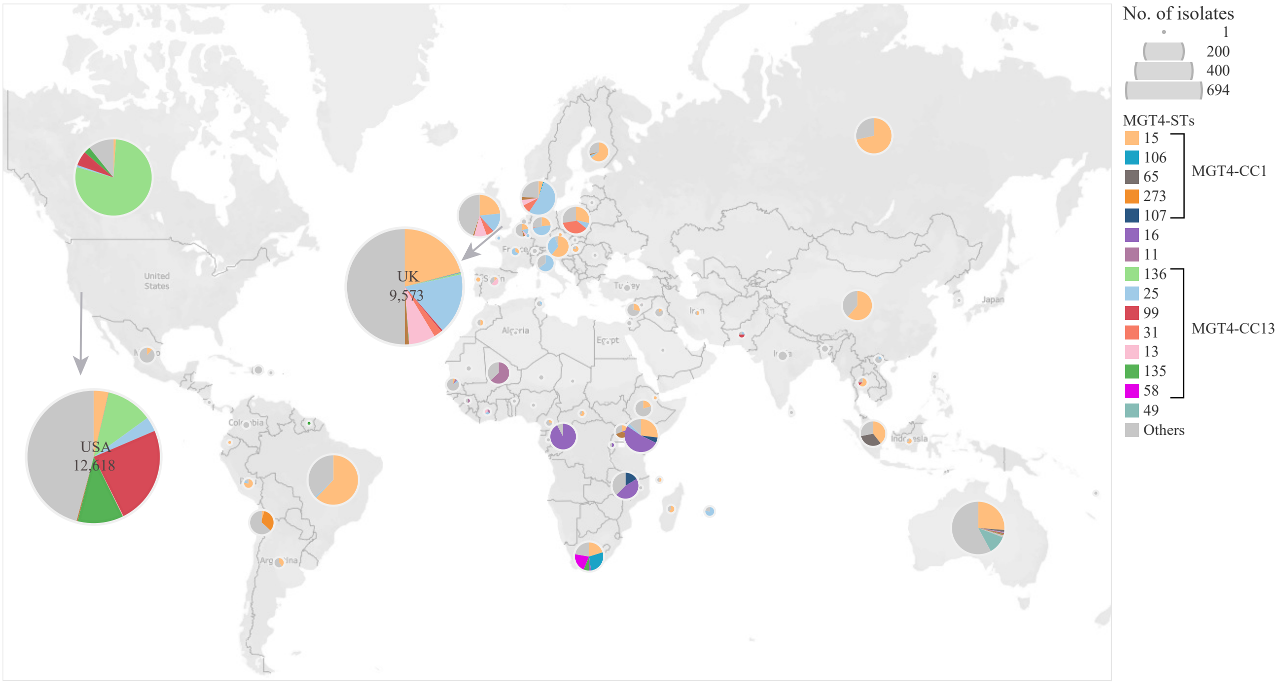


**Fig. S3. MGT4-STs distribution in different countries.** Although the majority of the *S.* Enteritidis genomes were from the USA and UK, there were 3017 isolates belonged to the other 86 countries. The top 15 MGT4-STs, representing 66.2% of the isolates, were represented with different colours. The size of each pie represented the number of isolates in each country except for USA (12,618) and UK (9,573). There were five top STs belonged MGT4-CC1 cluster, with MGT4-ST15 prevalent in most of the countries. Seven STs belonged to the MGT4-CC13 cluster, which were mainly observed in Canada and Europe countries. MGT4-ST16 was observed in Central/Eastern Africa, while MGT4-ST11 was prevalent in West Africa. MGT4-ST49 was unique to Australia and belonged to one of the two lineages outside the major global one. This map chart was created with Tableau v2019.2.

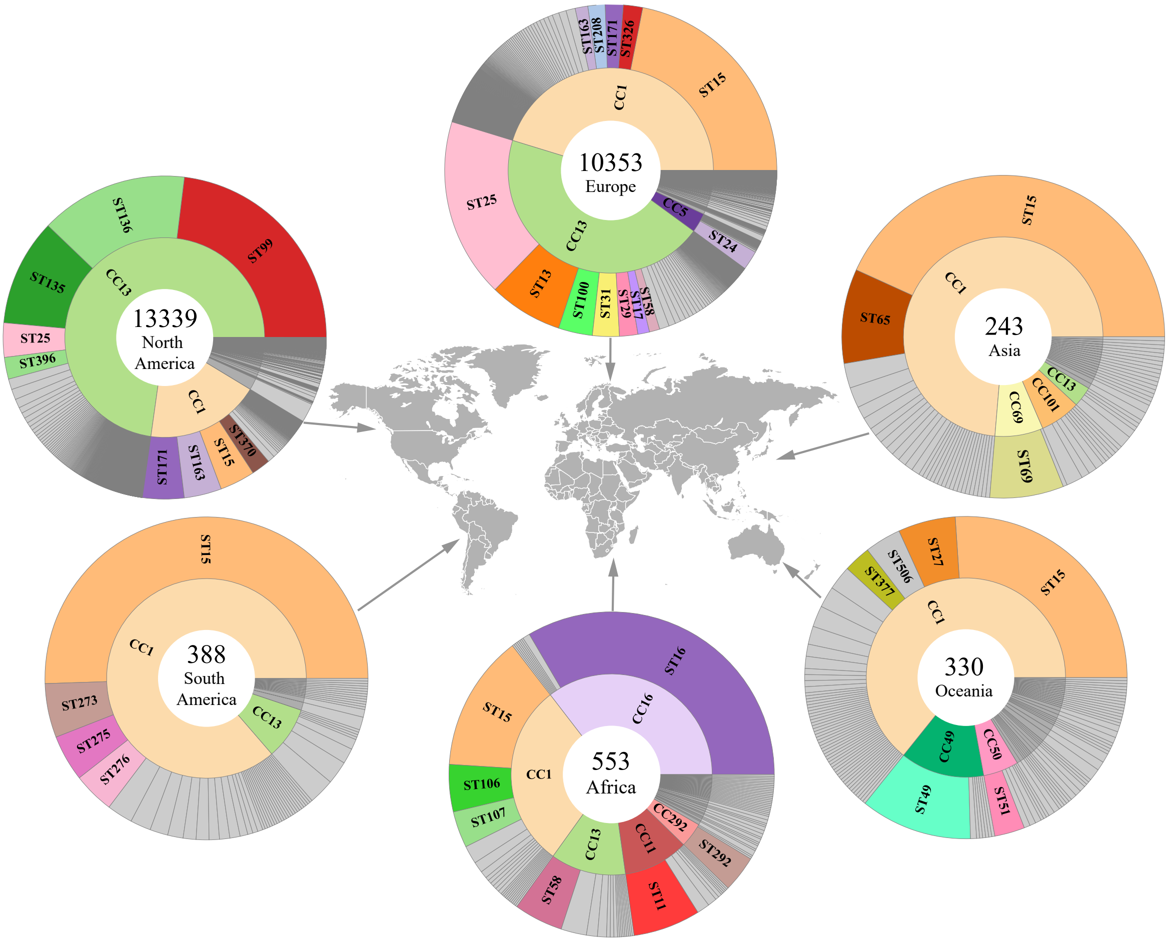


**Fig. S4. Continental restricted STs of *S.* Enteritidis were identified within each CCs at MGT4.** The numbers in each pie refer to the number of whole genome sequences analysed in each continent. The inner ring shows the main MGT4-CCs (with proportions of 3% or more) in each continent. MGT4-CC1 and CC13 were the two predominant MGT4-CCs with MGT4-CC1 being more widely distributed in all six continents. The outer ring shows the common MGT4-STs in each continent (with proportions of 2% or higher) within corresponding CCs. MGT4-ST15 was the dominant ST in MGT4-CC1, which were widely distributed all over the world. Continental restricted STs were identified, especially for the STs of MGT4-CC13, which was more prevalent in Europe and North America.

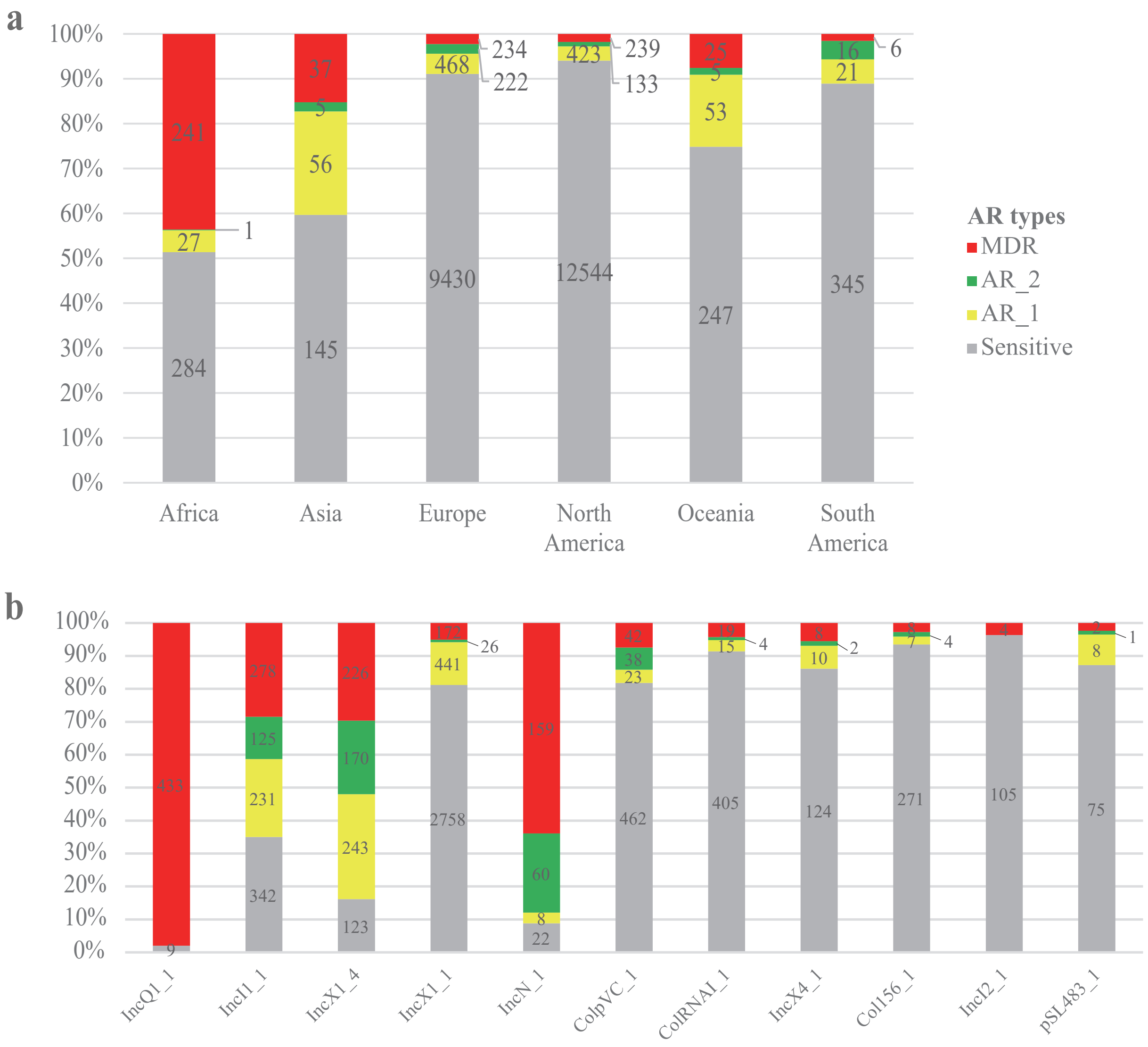


**Fig. S5. Geographic and predicted plasmid type distribution of MDR isolates. a.** Predicted antibiotic resistance of *S.* Enteritidis in different continents. The proportion of predicted antibiotic resistant isolates in different continents were shown with different colours representing different numbers of resistant drug classes. Isolates harbouring antimicrobial genes resistant to 3 or more drug classes (MDR) were represented with red colour, 2 different drug classes with green colour, 1 drug class with yellow colour, and no drug resistant gene with grey colour. **b.** Presence of AR genes in *S.* Enteritidis isolates with different predicted plasmid types. Different colors represent different types of AR. AR genes were of high proportion in *S.* Enteritidis isolates with IncQ1, IncN, IncI1, IncX1 plasmids. Particularly, 98.0% of the isolates with plasmid type IncQ1, were MDR associated.

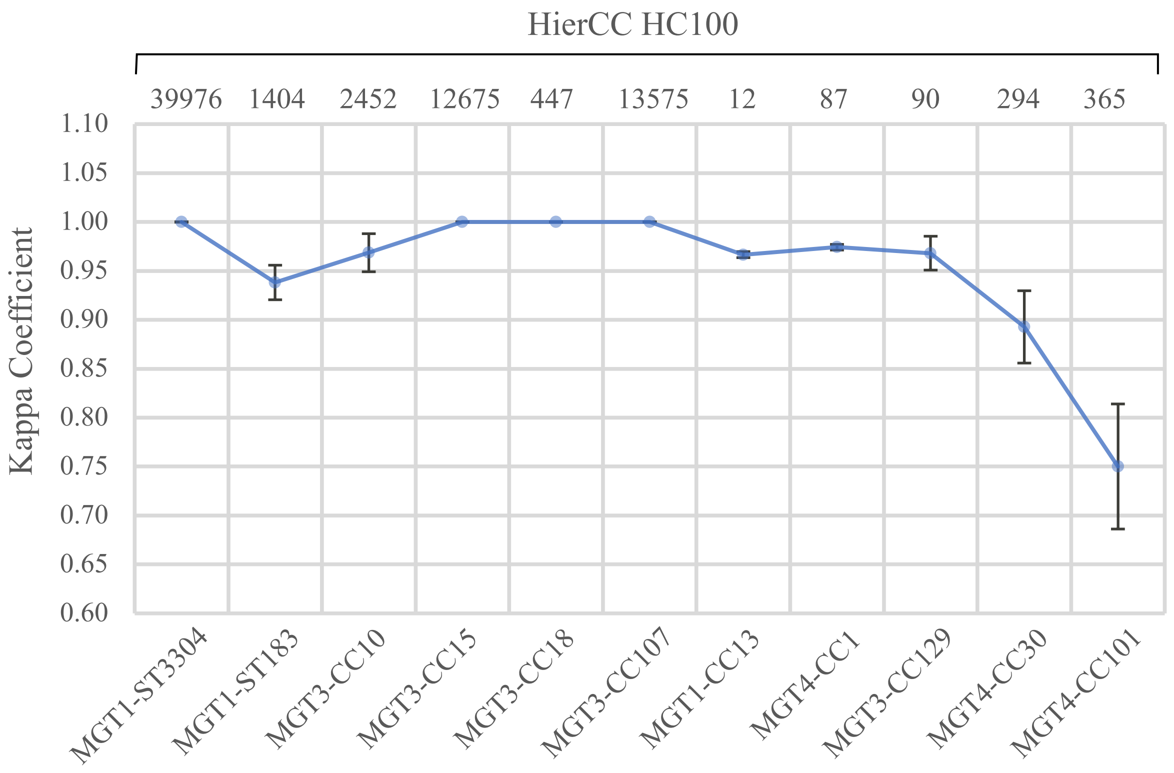


**Fig. S6. Population structure markers of *S.* Enteritidis by MGT STs/CCs and HierCC HC100 cluster types.** Three MGT1 (or 7 gene MLST) STs were representative of the Oceania prevalent clade A and clade C *S.* Enteritidis isolates, with MGT1-ST3304 as the main type. We identified 10 main lineages of clade B *S.* Enteritidis, which can be represented by 10 MGT-STs/CCs markers. By HierCC HC100, there were 10 different HC100 types corresponding to the 10 MGT lineage markers. The Cohen's kappa coefficient was calculated for each pair of the MGT-ST/CC lineage marker and the matched HC100 type using SAS v9.4. And the 95% confidential interval of the Cohen's kappa coefficient were shown. The closer the coefficient value were to one, the higher agreement between the two methods were.

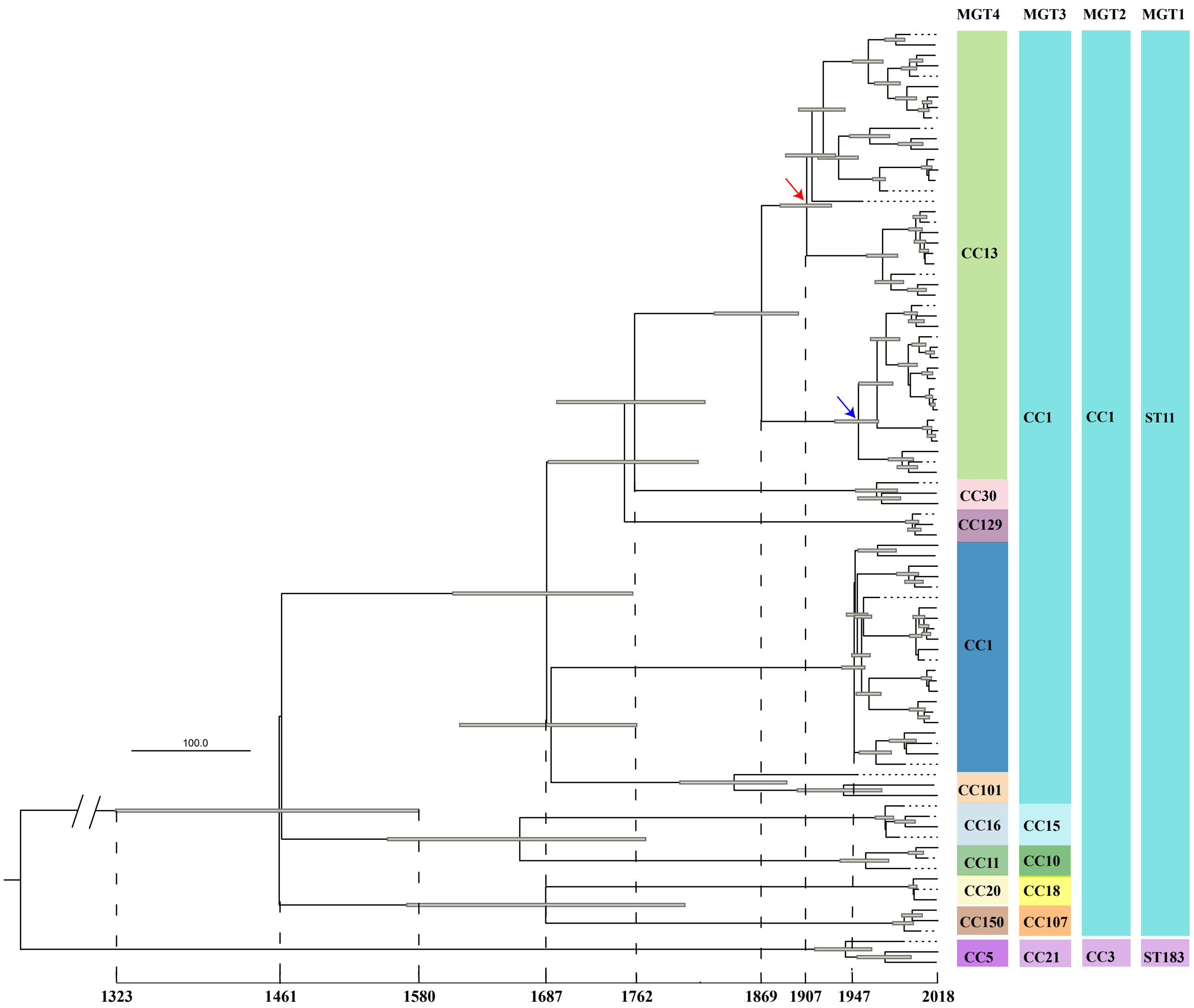


**Fig. S7. Evolutionary phylogeny of the global clade estimated with BEAST.** A total of 90 representative isolates were selected to generate a phylogeny with divergence date estimated using BEAST v1.10.4 with the uncorrelated relaxed log-normal clock and the Bayesian skyline model. The estimated divergence times were indicated for selected ancestral nodes with the 95% year confidence interval shown as error bars at each node. The CCs from MGT2 to MGT4 levels are shown next to the isolates, and the majority of the MGT2-CC1 isolates belonged to ST11 by 7 gene MLST (or MGT1). The most recent common ancestor (MRCA) of the 9 lineages belonging to MGT2-CC1, dated back to around 1460s (95% CI 1323-1580). Around 1687 (95% CI 1608-1760), MGT3-CC1 was separated into three lineages of MGT4-CC1, CC13 and CC101. Around 1869 (95% CI 1829-1900), the MGT4-CC13 lineage evolved into two sub-lineages of varied MGT4-STs, with one more prevalent in North America (labelled with red arrow) and the other one sub-lineage more prevalent in the Europe (labelled with blue arrow).

### Supplementary methods:

#### Core gene and core intergenic region definition of *S.* Enteritidis

The *S.* Enteritidis isolates from Enterobase (during May, 2018) were sub-sampled to identify representative isolates in three different ways depending on the size of the ribosomal sequence type (rST) [1]. For the isolates of the rSTs with fewer than 10 isolates each, all of the isolates were sampled. For the isolates of the rST with more than 10 and fewer than 350 isolates assigned, 10 isolates for each type were randomly sampled. For the isolates of the rST with more than 350 isolates (rST1425 and rST3888), 2% of the isolates for each type were sampled randomly while including isolates from as many countries and collection years as possible. A total of 1801 genomes, which were selected to represent 283 rSTs collected over 101 years (1917-2018) in 56 different countries.

The trimmed raw reads were then assembled using SPAdes v3.13.0 [2]. Prokka v1.12 was used to predict and annotate genes [3]. Roary v3.11.2 was used for the determination of core genes, which were defined as the genes present in 99% of the representative genomes. Core genes were identified as present in each genome following the nucleotide identity threshold >= 96% and alignment coverage threshold >= 99%. Those thresholds were assessed as the optimal identity by comparing the number of core genes generated with different identity from 90% to 100% (**Fig. S1**) [4]. Core genes with duplicates in the Roary output (paralogues) were removed. Core intergenic regions were defined using Piggy v3.11.2 based on the Roary outputs [5], and the nucleotide identity and coverage thresholds were identical to those of core genes.

#### Phylogenetic analysis

To define the major clades of *S.* Enteritidis globally, 1508 isolates were sampled based on the STs of MGT6 (random sampling of one isolate per ST with 3 or more isolates). A phylogenetic tree was constructed using ParSNP v1.2, which called core-genome SNPs [6, 7]. The branches with more than 10 isolates on the phylogenetic tree were collapsed together with iTOL v4 [8].

Average mutation rates were estimated for the global epidemic clade B of *S.* Enteritidis and the two dominant lineages, MGT4-CC1 and CC13. For the global epidemic clade B, a total of 90 isolates were randomly sampled based on MGT4-CCs and STs (covering the main STs for each CC). For the mutation rate of MGT4-CC1 or CC13, representative isolates were randomly sampled based on the higher resolution level MGT5 and collections years. For each MGT5-CC type, one isolate from each collection year were randomly sampled. SNPs derived from MGT9 alleles were then concatenated as a multi-alignment file and potential recombinant SNPs were removed using both Gubbins v2.0.0 and Recdetect v6.0 [9, 10]. Beast v1.10.4 was then used to estimate the mutation rate based on mutational SNPs [11]. A total of 24 combinations of clock and population size models were evaluated with the MCMC chain of 100 million states. Tracer v1.7.1 was used to identify the optimal model and to estimate population expansion over time [12]. For the predominant clade of *S.* Enteritidis including isolates of multiple lineages, uncorrelated relaxed lognormal clock combined with the Bayesian skyline model was chosen by the effective sample size (ESS) of both mean rate and joint probability of over 300. For the two dominant lineages of *S.* Enteritidis, MGT4-CC1 or CC13, the strict clock and Bayesian skyline models were found to be the optimal combination model by the ESS of mean rate and joint probability of over 100. The large size of each CC (more than 300 isolates for the Beast analysis) made it hard to reach 200 of ESS even for the optimal models.

**Supporting Results:**

**Potential repeat sequencing bias evaluation**

To evaluate any bias that may be caused by resequencing of the same strain or isolate, we identified all isolates of the same ST based on MGT9 and same metadata based on collection country, collection year and month, and source type. Such isolates were conservatively treated as repeat sequencing of the same isolate and the "duplicates" were removed from the dataset. A total of 4,026 out of the 26,670 isolates (17.8%,) were identified and "duplicates" removed from the dataset. The prevalence of STs in different geographic regions, collection time and sources, were re-evaluated in the reduced dataset of 22,644 isolates and compared with the original dataset. Based on the number of isolates after resampling, the rank of STs and CCs in different geographic regions, collection time and sources were compared against the original dataset.

1. For the continental distribution of *S.* Enteritidis (compared against **Fig. 2)**:

   The global distribution of *S.* Enteritidis were reevaluated based on MGT4 STs and CCs. The top 10 MGT4 CCs (with >= 10 isolates each) in each continent remain the same after resampling, except for MGT4 CC378, which dropped from rank 9 to rank 12 in Europe (**Data Set S2, Tab 1**). To compare the rank of the CCs before and after sampling, Kendall's tau value was calculated [13]. If Kendall's tau value is close to 1, it indicates strong agreement. If Kendall's tau value is close to -1, it indicates strong disagreement. The ranks of CCs in each continent before and after sampling strongly agreed, with Kendall's tau value of 0.917 (P < 0.001).

   The top 20 MGT4 STs (with >= 10 isolates each) in each continent were also the same, except for MGT4-ST2060 dropped from rank 15 to 24 in North America (**Data Set S2, Tab 2**). The ranks of STs agreed strongly with Kendall's tau of 0.944 (P < 0.001).

2. For the states and source distribution of *S.* Enteritidis in the USA (compared against **Fig. 4**):

   The top 10 MGT4 STs in the USA remained the same before and after the resampling (**Data Set S2, Tab 3**). The rank of the STs across all states has a Kendall's tau value as 0.987 (P < 0.001).

3. For the temporal analysis of *S.* Enteritidis in UK (compared against **Fig. 5**):

   For the rank of UK MGT5 STs in different months before and after "duplicates" removal, the top 13 MGT5 STs remain the same (**Data Set S2, Tab 4**). The Kendall's tau value based on the rank of STs in each month was 0.900 (P value < 0.001).

4. For the outbreak propensity evaluation between MGT4-CC1 and CC13 using the clinical *S.* Enteritidis isolates from UK during 2014 to 2018 (compared against **Data Set S1, Tab 4 and Tab 5**):

The top 12 STs for large outbreak clusters evaluation remain the same (**Data Set S2, Tab 5).** Although the predicted OR value adjusted slightly, the P values remain significant. It must be noted that the artificial removal of "duplicates" effectively removed many potential outbreak isolates. Importantly, MGT4-CC13 was confirmed to be significantly associated with large outbreak clusters than MGT4-CC1 (**Data Set S2, Tab 6**).

In summary, by removing potential "duplicates", recalculation and comparison with the original dataset, no significant difference was observed in either top STs and their ranks. Thus, any bias brought by potential repeat sequencing of the same isolates or even same strain from an outbreak was negligible.
